# Model-Based Meta-Analysis of Relapsing Mouse Model Studies from the Critical Path to Tuberculosis Drug Regimens Initiative Database

**DOI:** 10.1101/2021.09.13.460195

**Authors:** Alexander Berg, James Clary, Debra Hanna, Eric Nuermberger, Anne Lenaerts, Nicole Ammerman, Michelle Ramey, Dan Hartley, David Hermann

## Abstract

Tuberculosis (TB), the disease caused by *Mycobacterium tuberculosis* (*Mtb*), remains a leading infectious disease-related cause of death worldwide, necessitating the development of new and improved treatment regimens. Non-clinical evaluation of candidate drug combinations via the relapsing mouse model (RMM) is an important step in regimen development, through which candidate regimens that provide the greatest decrease in probability of relapse following treatment in mice may be identified for further development. Although RMM studies are a critical tool to evaluate regimen efficacy, making comprehensive “apples to apples” comparisons of regimen performance in the RMM has been a challenge, in large part due to the need to evaluate and adjust for variability across studies arising from differences in design and execution. To address this knowledge gap, we performed a model-based meta-analysis on data for 17 unique regimens obtained from a total of 1592 mice across 28 RMM studies. Specifically, a mixed-effects logistic regression model was developed that described the treatment duration-dependent probability of relapse for each regimen and identified relevant covariates contributing to inter-study variability. Using the model, covariate-normalized metrics of interest, namely treatment duration required to reach 50% and 10% relapse probability, were derived and used to compare relative regimen performance. Overall, the model-based meta-analysis approach presented herein enables cross-study comparison of efficacy in the RMM, and provides a framework whereby data from emerging studies may be analyzed in the context of historical data to aid in selecting candidate drug combinations for clinical evaluation as TB drug regimens.

## INTRODUCTION

*Mycobacterium tuberculosis* (*Mtb*), the causative agent of tuberculosis (TB), infects an estimated one quarter of the world’s population, and causes an estimated 1.4 million TB-related deaths per year, making it the leading worldwide cause of death due to a single infectious disease excluding severe acute respiratory syndrome coronavirus 2 (SARS-CoV-2) (1–3). Despite a few successes in the past decade in bringing forward new treatments for drug-resistant pulmonary TB (4–6), progress in advancing new drugs and regimens for pulmonary TB has been limited. The standard of care treatment regimen for drug-susceptible pulmonary TB, based on the combination of isoniazid, rifampin, pyrazinamide, and ethambutol, remains essentially unchanged for more than 30 years (7). While the regimen is efficacious (∼90% cure rates in a clinical trial setting), its complexity, long treatment duration, often poor tolerability, and requirement for high adherence to minimize treatment failure represent gaps that must be addressed through the development of novel treatment regimens. Treatment with the standard regimen is typically six to nine months. The major aim of ongoing TB drug research is to develop markedly shorter treatments (e.g., less than two months) to improve adherence and overall cure rates.

To accelerate progress in fighting the global TB pandemic, organizations such as the Bill and Melinda Gates Foundation have worked to promote renewed interest in TB drug development through funding of collaborative efforts to accelerate the development of new drug regimens. One such effort, the Critical Path to TB Drug Regimens (CPTR) Initiative, was formed to facilitate the development of new tools and methodologies for use in TB drug and regimen development (8). Since its inception, a primary focus of the CPTR Initiative has been the aggregation and standardization of clinical and non-clinical datasets to enable pooled analyses of existing data on TB drug regimens. These analyses, such as the TB-ReFLECT meta-analysis (9), provide greater insight into questions underlying the research and development of new drugs and regimens to guide the design of new studies and selection of regimens for further evaluation.

The recent uptick in investment and collaborative efforts in TB research and development has resulted in the identification of numerous drug candidates with potential for efficacy in the treatment of both drug-susceptible and drug-resistant TB. Given the multitude of potential combinations of existing and novel drugs, the prioritization of candidate regimens is now a significant challenge, especially for candidate regimens for which clinical data are not yet available for one or more regimen components. The selection of such novel regimens for further advancement into clinical studies relies heavily on the comparison of non-clinical efficacy studies (10), particularly the relative performance of regimens in achieving non-relapsing cure. This endpoint is commonly assessed using the relapsing mouse model (RMM) (11), a murine model of TB which tests the overall curative potential of a drug combination by evaluating the proportion of mice exhibiting relapse following different treatment durations. Relapse is defined as recurrence of *Mtb* growth in cultures of lung (and sometimes spleen) tissue samples obtained post-sacrifice after a post-treatment clearance period of typically three to six months. Although this definition of relapse implies the occurrence of no culture growth at the end of treatment, operationally this is not an absolute requirement to determine the treatment duration required to prevent relapse. Comparison of regimens in the RMM is typically done by rank-ordering based on the overall proportion of relapse at various treatment durations. Depending on the effect size, regimens that exhibit lower proportions of relapse following the completion of treatment and/or similar proportions of relapse following shorter treatment durations may be considered as potential improvements upon standard of care regimens and thus may be considered for clinical evaluation.

Although RMM studies are a critical tool in comparing regimen performance, the model has limitations with regard to study duration, design heterogeneity, and overall utility for decision-making. Designs can vary widely from lab to lab, with differences across multiple design elements such as mouse strain, bacterial strain, inoculation dose and route, recovery duration and bacterial culture methods. These inter-study differences, not to mention regimen-specific differences in treatment duration and dose selection, contribute to the observation that the efficacy of a specific regimen in the RMM can vary widely from study to study, even within the same lab, confounding decision-making (11–13). This inter-study variability obviates the comparison of regimens across studies, which is a key limitation when attempting to prioritize regimens as logistical considerations typically limit RMM studies to the evaluation of only a limited number of regimens in a given study. Moreover, the analysis of regimen performance within a given study has historically utilized conventional group-to-group statistical comparisons designed to evaluate the proportion of mice relapsing at a relatively small number of treatment durations (e.g., three to five). Lenaerts, et al., have demonstrated that the RMM requires at least 15 animals per regimen to detect a 50% reduction in relapse probability at a given treatment duration with at least 80% power (14). This analysis strategy based upon point-by-point comparison of relapsing proportions limits the utility of the model as the differences between candidate regimens are often much smaller. Further, the focus on comparative efficacy at pre-specified treatment durations requires careful selection of multiple treatment durations (and larger numbers of mice) to be tested to ensure identification of regimens that achieve significant rates of cure (e.g., 10% or lower probability of relapse) with a shorter treatment duration than the current standard of care regimen.

To improve the overall utility of RMM study data and interpretation for regimen prioritization, the CPTR Initiative applied modeling and simulation-based techniques to the analysis of RMM study data across multiple pooled, historical datasets. The objective of this work, which was based on data from a total of 28 studies conducted in two separate laboratories, was to develop a suitable model-based regression approach to: 1) determine the treatment duration-dependent relapse probability profiles for the included drug regimens, 2) quantify the magnitude of inter-study variability in treatment response and assess the impact of study-level covariates on treatment response, and 3) calculate key metrics of interest, namely treatment duration required to reach 50% and 10% relapse probability (T_50_ and T_10_, respectively), for comparison of relative regimen performance in the RMM when adjusted for study-level covariates. By applying a model-based analysis, it was expected that inter-study differences could be accounted for and unbiased estimates of informative parameters, such as T_10_, could be reported to support decision-making. Importantly, the model-based approach was also expected to yield uncertainty in parameters, such as T_10_, which will be a function of the amount of and consistency in available data, further contributing to informed decision-making.

## METHODS

### Data

The analysis was conducted based on data from studies conducted by the laboratories of Dr. Anne Lenaerts (Colorado State University, CSU), and Dr. Jacques Grosset and Dr. Eric Nuermberger (Johns Hopkins University, JHU). The experimental details of the contributed datasets have been described elsewhere (13, 15–23). Datasets corresponding to the 25 studies were standardized to a data template developed by the authors as part of the CPTR Initiative prior to aggregation into the pooled analysis dataset for initial model development. Data from three additional JHU studies were subsequently added to the pooled dataset, resulting in a total of 28 studies in the final dataset (Table S-1). All studies were performed in BALB/c mice inoculated via aerosol inhalation with either *Mtb* Erdman TMC 107 (ATCC 35801) (CSU) or mouse-passaged H37Rv (ATCC 27294) (JHU) *Mtb* strains. In all studies, the primary endpoint for each mouse was relapse as a binary (0/1) variable, determined on the basis of whether *Mtb* growth was present or absent upon culturing a homogenate of lung tissue on solid media. Data from untreated control animals were excluded from the analysis.

A total of 17 unique regimens were available for analysis including the following drugs and doses: bedaquiline (B) 25 mg/kg, ethambutol (E) 100 mg/kg, isoniazid (H) 10 mg/kg or 25 mg/kg, linezolid (L) 100 mg/kg, moxifloxacin (M) 100 mg/kg, rifampin (R) 10 mg/kg, pretomanid (Pa) 100 mg/kg, and pyrazinamide (Z) 150 mg/kg. Treatments were administered orally once per day on a five out of seven days per week dosing schedule in all studies, with the exception of one study which dosed moxifloxacin 100 mg/kg twice daily for a total daily dose of 200 mg/kg. Regimen designations were assigned based on standard abbreviations for regimen components, with regimens containing an initial two-month (8-week) intensive phase followed by a separate continuation phase delineated by a “/” according to convention (i.e., HRZE/HR). Treatment duration in months was included as the independent, continuous variable for the analysis.

Study-level covariates included in the analysis dataset included categorical variables (i.e., 12-week versus 24-week recovery duration post-treatment, type of solid culture media used to assess *Mtb* growth, fixed versus time-variable dosing, number of inoculation groups) and continuous variables (i.e., culture incubation period [days], inoculum size for aerosol [log_10_ CFU], infection duration prior to treatment [days], lower limit of detection for bacterial growth [log_10_ CFU], and average baseline lung bacterial burden as determined in control animals at start of treatment [log_10_ CFU]). An additional categorical “Lab” covariate designating the contributing laboratory was included due to the high degree of within-lab correlation across studies of the following variables, which were excluded from analysis: average mouse age, *Mtb* strain, and *Mtb* cultivation method for inoculum preparation. Data imputation was not performed except for one study missing the inoculum size and which was therefore imputed as the median value from the dataset.

### Model development

The model presented herein was developed in two stages, with the first stage representing primary model development based on the initial dataset of 25 studies, which was subsequently followed by a second stage whereby the model was updated based on data from three additional studies.

During each stage, exploratory data analysis was performed to evaluate the informational content of the dataset and assess consistency across studies. Independent, dependent, and treatment variables were explored as well as relevant covariate information. Graphical outputs were used to help determine significant trends in covariates in order to aid in the model-building process.

Model-based analysis was performed using a nonlinear mixed-effects modeling approach. Specifically, a logistic regression model was developed with relapse treated as a binary 0 or 1 endpoint corresponding to absence or presence of relapse, respectively, treatment duration as an independent variable, and study as a random effect. Determination of model performance during development and refinement was on the basis of standard goodness-of-fit plots and summary statistics, as well as evaluation of log likelihood-based metrics (i.e., objective function value [OFV] and Akaike’s Information Criterion [AIC]) (24). The general model structure is described by Equations 1 through 3:

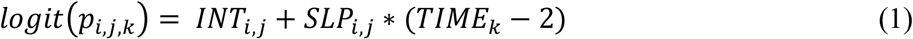

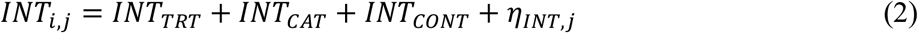

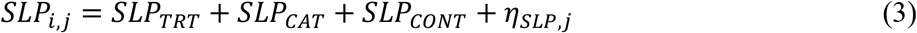

Where:

- *p*_*i,j,k*_ is the probability of relapse for a given treatment/covariate combination *i* in Study *j* at treatment duration *k*
- *INT*_*i,j*_ is the logit relapse probability after two months of treatment for the *i*th regimen in the *j*th study with a study-level covariate effect
- *SLP*_*i,j*_ is the slope of the logit relapse probability versus treatment duration for the *i*th regimen in the *j*th study with a study-level covariate effect
- TRT is a categorical treatment indicator
- CAT denotes a categorical covariate effect
- CONT denotes a continuous covariate effect
- η is the random effect of the *j*th study for intercept and slope, assumed to be N(0,σ^2^)

Metrics of interest, namely T_10_ and T_50_, were calculated as follows from the model estimates according to Equations 4 and 5:

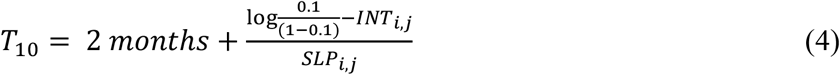

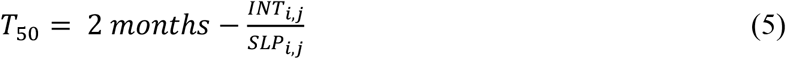

Treatment-specific intercept (INT) and slope (SLP) values (viz., *INT*_*i*_ and *SLP*_*i*_) were estimated where supported. Regimens with a two-month intensive phase that was identical to another regimen (e.g., HRZE/HR and HRZE) were modeled as having the same INT value. Regimens with similar components were grouped together for INT, SLP, or both parameters when necessary to maintain model stability while maximizing the ability to differentiate between regimens.

Covariates were assessed for significance on INT and SLP terms using graphical and stepwise covariate modeling during the first stage of the analysis. Stepwise covariate modeling using the likelihood ratio test was performed using respective alpha levels of 0.05 and 0.01 for forward and backward phases. All covariates were evaluated through the inclusion of additive fixed effects on either INT or SLP. Fixed effects for continuous covariates represent the estimated effect of the covariate at the dataset median value as all continuous covariates were centered for analysis. Covariate effects were re-evaluated during the second stage of the analysis to confirm that the inclusion of additional data did not warrant a change in the model structure.

### Model evaluation

Simulation-based diagnostics (i.e., visual predictive checks [VPCs]) were performed on pivotal models to evaluate model predictive performance when stratified by regimens and covariates. A total of 1000 replicates were simulated and 95th percentiles of the simulated values were overlaid with the observed values to visually assess the agreement between the model-based treatment duration-dependent relapse probabilities and the observed relapse proportions at the various treatment durations.

To obtain estimates of model precision and support comparison of the various regimens, a non-parametric bootstrap approach was employed. A total of 500 replicates of the analysis dataset were generated with replacement with stratification by regimen. Model parameters were obtained through re-estimation of the model on each replicated dataset, with T_10_ and T_50_ estimates generated from the bootstrap model estimates at the covariate reference value to generate covariate-normalized confidence intervals. Rank orders were calculated across all regimens using the median T_10_ and T_50_ values from the bootstrap runs.

### Simulations

Simulations were run to examine the effects of covariates on T_10_ and T_50_. Observed inoculum values in the source dataset were used to generate prediction curves from the model estimates for HRZE/HR at the median baseline bacterial burden value. This was repeated for the observed baseline bacterial burden at the median inoculum value, as well as for all combinations of inoculum and baseline bacterial burden values in the dataset.

### Software and hardware

All data assembly and analysis was performed in R (25) as implemented via the RStudio environment (26). Logistic regression was done using the *glm* function in the stats package. Diagnostic metrics and plots were generated via the *dx* function in the *LogisticDX* (27) package and using the *ggplot2* (28) package.

## RESULTS

### Exploratory data analysis

Data from 1310 mice were included in the first stage of the analysis of 25 studies, with data from three more studies contributing an additional 282 mice added in the second stage, for a grand total of 1592 mice across 28 studies. Summary statistics across all studies in the analysis are shown in Table 1, with a further summary of data for each regimen provided in Table 2.

**TABLE 1.**
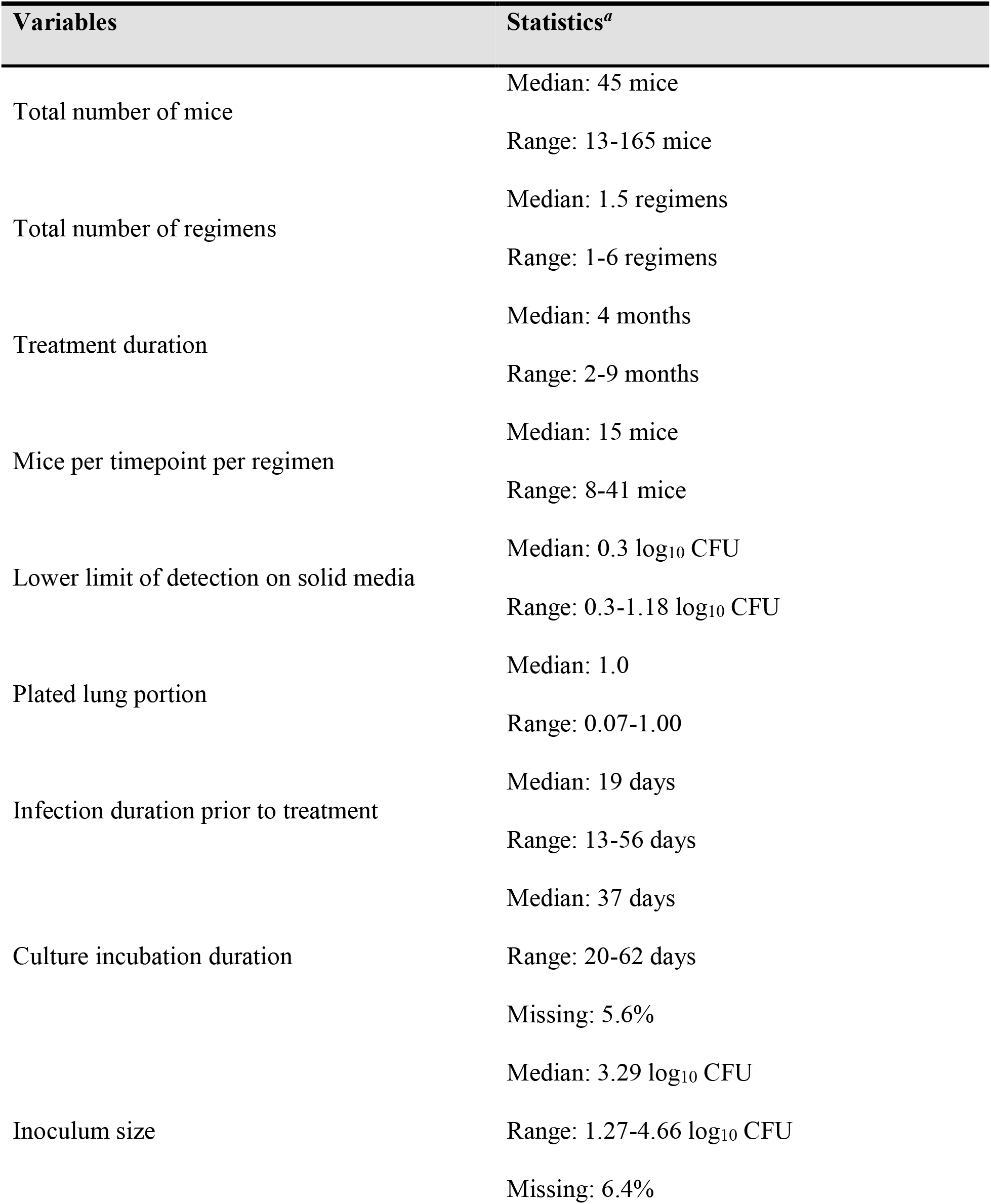
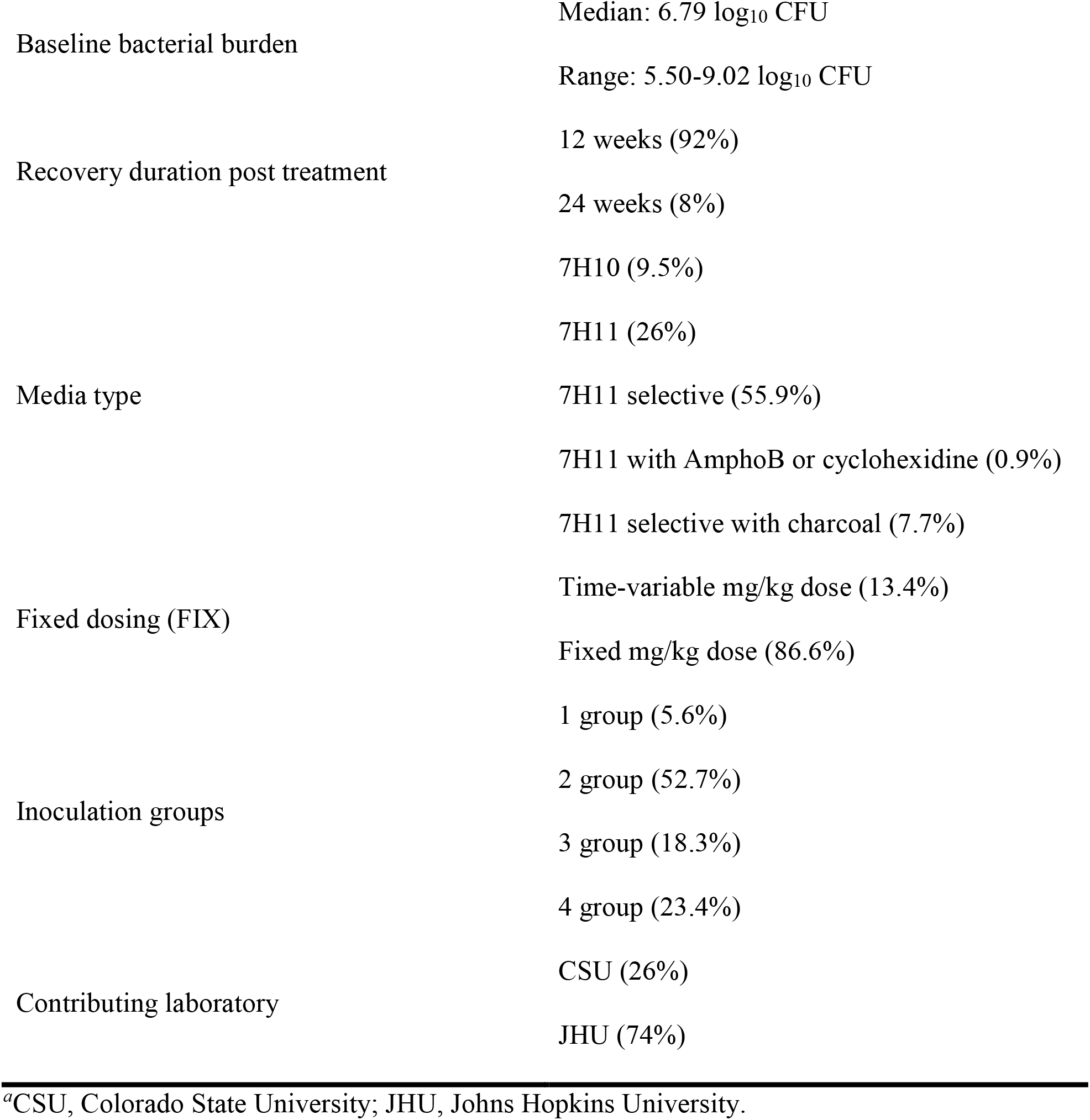
Summary statistics for study-level data

**TABLE 2.** Summary of available data by regimen

| Regimen<sup>a</sup> | Number of studies | Number of mice | Assessed treatment durations (months) |
| --- | --- | --- | --- |
| BPa | 3 | 79 | 2, 3, 4 |
| BPaL | 4 | 139 | 2, 3, 4 |
| HE | 1 | 30 | 6, 9 |
| HRE | 1 | 31 | 3, 6 |
| HRE/HR | 1 | 45 | 3, 4.5, 6 |
| HRZ/HR | 15 | 531 | 2, 2.5, 3, 4, 5, 6 |
| HRZE | 5 | 105 | 2, 3, 4 |
| HRZE/HR | 9 | 199 | 3, 4, 4.5, 5, 6 |
| HRZE/HRZ | 1 | 15 | 3 |
| HRZM | 2 | 40 | 2, 3 |
| HRZM/HRM | 2 | 23 | 3, 6 |
| HZE | 1 | 29 | 3, 6 |
| RMZ | 1 | 36 | 3, 4, 5 |
| RMZ/RM | 5 | 215 | 3, 4, 5, 6 |
| RMZ/RM<sup>b</sup> | 1 | 15 | 2.5 |
| RMZE | 2 | 45 | 2, 3 |
| RMZE/RM | 1 | 15 | 3 |

<sup>a</sup>B, bedaquiline; E, ethambutol; H, isoniazid; L, linezolid; M, moxifloxacin; Pa, pretomanid; R, rifampin; Z, pyrazinamide.
<sup>b</sup>Moxifloxacin 100 mg/kg dosed twice daily for a total daily dose of 200 mg/kg.

Observed relapse proportions for each regimen by treatment duration are shown in Fig. 1, with results grouped by study and stratified by contributing laboratory. Significant variability in response was observed across studies, which was clearly visible during exploratory analysis for the most commonly utilized control regimen, HRZ/HR. As judged by graphic depiction of raw data, the HRZ/HR treatment duration to reach 50% relapse probability (i.e., 50% proportion of mice exhibiting relapse) ranged from less than 2.5 months to greater than 4.5 months. This observation is consistent with previous observations of variable treatment response for regimens across studies by using different mouse models (12), and is expected given the differences in study design and covariates. To quantify the magnitude of the inter-study variability and investigate which covariates may be potential sources of variability, a mixed-effects logistic regression modeling approach was applied under the assumption of logit-linearity (which was confirmed prior to analysis; refer to Fig. S-1).

**FIG 1.**
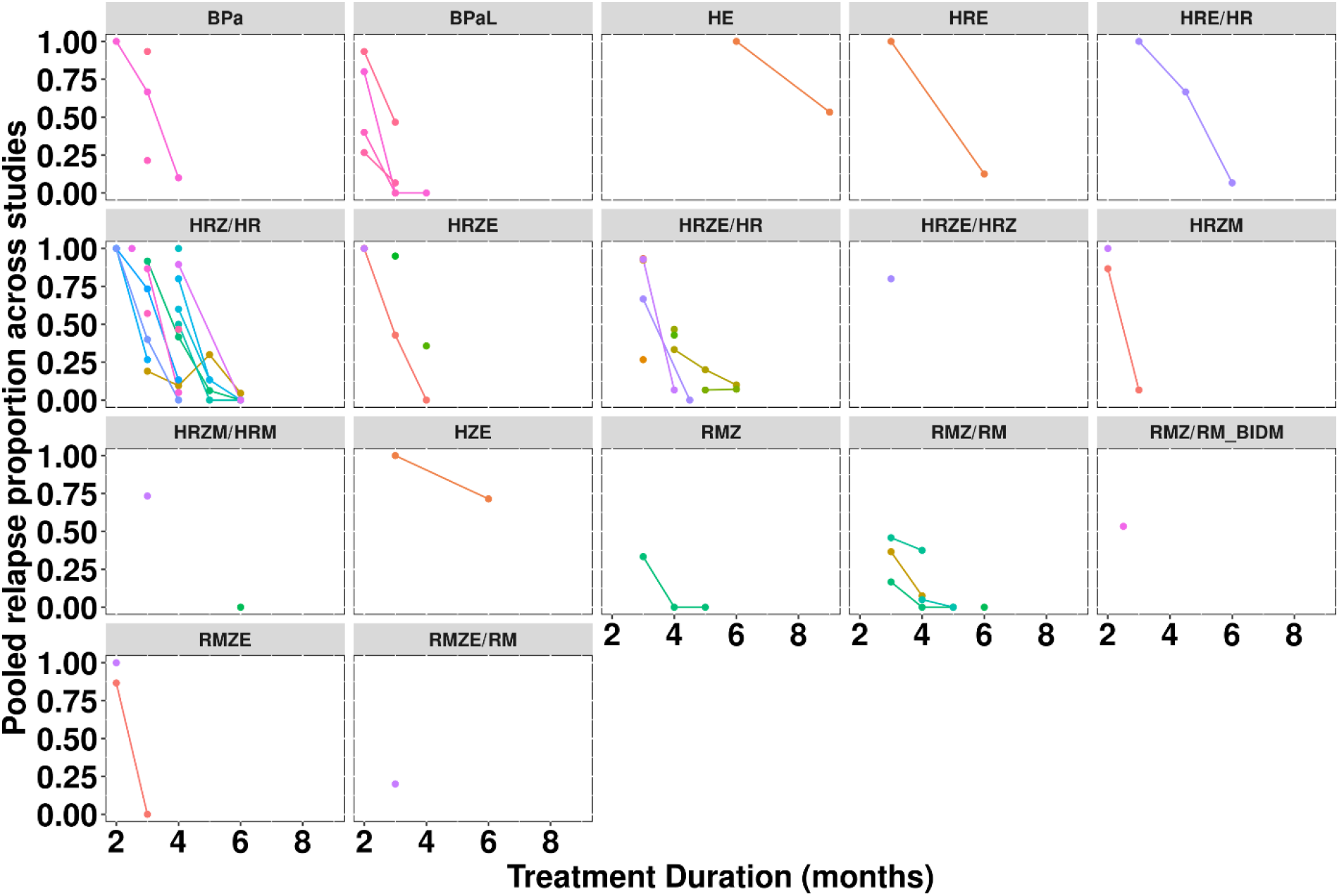
Relapse proportion by treatment duration across stratified by regimen and study. Each study is grouped together with lines representing the relapse-time course in a particular study. Points represent individual timepoints in a particular study. “RMZ/RM_BIDM” denotes a version of the RMZ/RM regimen where moxifloxacin was administered at 100 mg/kg twice daily for a total daily dose of 200 mg/kg. M, moxifloxacin; R, rifampin; Z, pyrazinamide.

### Model development and evaluation

Key steps in model development are summarized in Table 3, which lists the pivotal model runs from the primary analysis stage. As expected, inclusion of treatment-specific INT and SLP terms greatly improved the model fit as compared to a “naive” model including shared INT and SLP terms across all treatments and study as a random effect on both INT and SLP. Subsequent model simplification to group similar treatments together to improve model stability prior to covariate analysis further reduced the AIC value of the model. Stepwise covariate analysis resulted in the identification of inoculum amount (INOC) and average baseline lung bacterial burden (BASE) as significant covariates on INT and SLP, respectively. This further reduced both the OFV and the AIC values, and eliminated trends in goodness-of-fit plots of random effects (η_j_ values) versus covariates (not shown). Further, the inclusion of these covariates accounted for a significant amount of inter-study variability, decreasing the inter-study standard deviation estimates for INT (σ_INT_) and SLP (σ_SLP_) by 37% and 23%, respectively. Subsequent model refinements during the primary analysis phase focused on separating INT and/or SLP terms for treatment regimens that were grouped together prior to covariate analysis while maintaining model stability. This was done to support our primary objective of extracting treatment duration-dependent relapse probability profiles from the model to assess differences in response between regimens, and resulted in further model improvements. In the second stage of the analysis, the model was updated to support the inclusion of additional datasets that became available during the course of the project. This update included the addition of treatment-specific INT and SLP terms for the BPa and BPaL regimens not included in the original dataset, but did not result in any further modification of the model structure. Goodness-of-fit plots for the final model are presented in Fig. S-2. The final parameter-covariate relationships are shown below:

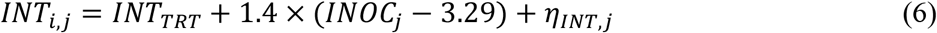

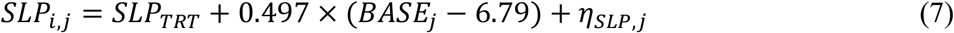

**TABLE 3.** Listing of pivotal models during primary analysis phase

| Model number | Description | OFV<sup>a</sup> | AIC<sup>b</sup> |
| --- | --- | --- | --- |
| 3 | Naïve model | 1131.2 | 1141.2 |
| 25 | “Full” treatment effect, no covariates | 987.1 | 1033.1 |
| 35 | “Reduced” treatment effects, no covariates | 990.1 | 1018.1 |
| 37 | Reduced treatment effects plus covariates | 899.2 | 931.2 |
| 44 | Final model | 882.4 | 920.4 |

<sup>a</sup>OFV, objective function value.
<sup>b</sup>AIC, Akaike’s Information Criterion.

Estimates for the treatment-specific fixed-effects parameters are presented in Table 4, whereas random-effects estimates from the final model were 1.209 and 0.636 for σ_INT_ and σ_SLP_, respectively, with an estimated correlation of –0.75.

**TABLE 4.** Treatment-specific fixed-effects parameter estimates

| Regimen<sup>a</sup> | INT<sub>TRT</sub> |  | SLP<sub>TRT</sub> |  |
| --- | --- | --- | --- | --- |
|  | Estimate | Standard error | Estimate | Standard error |
| BP<sub>a</sub> | 2.27 | 1.11 | -1.96 | 0.892 |
| BP<sub>a</sub>L | 0.499 | 0.78 | -3.77 | 0.782 |
| HE | 21.5 | 2.42 | -3.00 | 0.27 |
| HRE or HRE/HR | 9.58 | 0.998 | -3.00 | 0.27 |
| HRZ/HR | 4.86 | 0.481 | -3.00 | 0.27 |
| HRZE or HRZE/HRZ | 4.7 | 0.438 | -3.23 | 0.317 |
| HRZE/HR | 4.7 | 0.438 | -3.00 | 0.27 |
| HRZM | 3.61 | 0.759 | -4.79 | 1.26 |
| HRZM/HRM | 3.61 | 0.759 | -3.00 | 0.27 |
| HZE | 12.7 | 1.33 | -3.00 | 0.27 |
| RMZ/RM_BIDM<sup>b</sup> | 0.654 | 1.16 | -3.11 | 0.492 |
| RMZ | 2.22 | 0.885 | -3.54 | 0.787 |
| RMZ/RM | 2.22 | 0.885 | -3.11 | 0.492 |
| RMZE | 2.11 | 0.771 | -3.54 | 0.787 |
| RMZE/RM | 2.11 | 0.771 | -3.11 | 0.492 |

<sup>a</sup>B, bedaquiline; E, ethambutol; H, isoniazid; L, linezolid; M, moxifloxacin; Pa, pretomanid; R, rifampin; Z, pyrazinamide.
<sup>b</sup>Moxifloxacin 100 mg/kg dosed twice daily for a total daily dose of 200 mg/kg.

To assess predictive capability of the final model, a VPC was performed with stratification by treatment (Fig. 2). Observed relapse proportions were generally within the 95% prediction interval for the various regimens, although underprediction was observed at a single treatment duration for a few regimens. Additional VPCs with stratification by study-specific covariate values for INOC and BASE as well as by study (Fig. S-3 to Fig. S-5) show a similar pattern of agreement between observed and predicted values. Taken together, the VPCs indicate that the model exhibits acceptable predictive performance.

**FIG 2.**
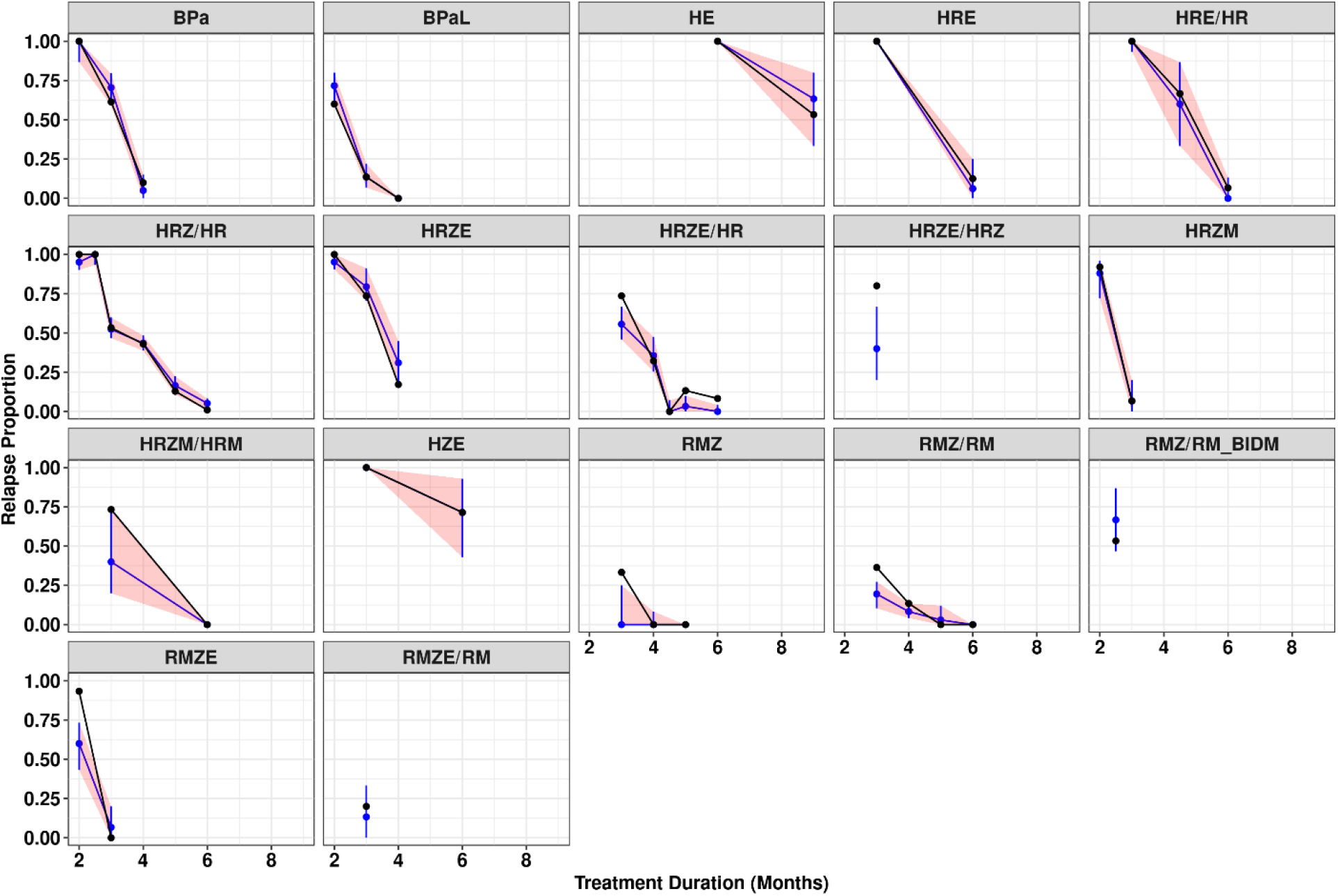
Visual predictive check for the final model – stratified by regimen. Black dots and solid lines represent the observed relapse proportion. Blue lines represent the median prediction from the final model. The red shaded area represents the 90% prediction interval. “RMZ/RM_BIDM” denotes a version of the RMZ/RM regimen where moxifloxacin was administered at 100 mg/kg twice daily for a total daily dose of 200 mg/kg. M, moxifloxacin; R, rifampin; Z, pyrazinamide.

### Comparison of regimen efficacy

A bootstrap analysis was performed to compare the efficacy of the various treatment regimens and obtain distribution-independent precision estimates. Relapse probability versus treatment duration profiles, derived at the covariate median values using estimates obtained from each bootstrap replicate, are shown in Fig. 3 along with HRZE/HR included for reference as the clinical standard of care regimen. Covariate-normalized T_50_ and T_10_ values were also obtained from each bootstrap replicate and are depicted in Fig. 4 (Panels A and B, respectively), with median and 95% confidence interval (CI) values provided in Table 5 along with regimen rank order based on the median values. As seen in Fig. 3, efficacy profiles are generally well estimated for most treatment regimens, although BPa exhibits a relatively large confidence interval attributed to poorer precision in the BPa-specific SLP parameter secondary to the relatively small number of treatment durations and mice available for this regimen. This is consistent with the forest plots, which show that the T_10_ confidence interval for BPa is much wider for this regimen as compared to the other regimens. Although T_10_ generally shows a slightly broader confidence interval for all regimens as compared to T_50_, both parameters show the same three groupings based upon whether their confidence interval overlapped the median estimate for HRZE/HR: 1) regimens with better performance (HRZM, RMZ/RM, RMZ, RMZE/RM, RMZE, and BPaL), 2) regimens with similar performance (HRZ/HR, HRZE, HRZE/HRZ, BPa, and HRZM/HRM); and 3) regimens with poorer performance (HE, HRE, HRE/HR, and HZE). Similarly, rank ordering demonstrated that BPaL was the best performing regimen for both T_10_ and T_50_ values (median values of 2.13 months and 2.69 months, respectively), with ranks for other regimens also consistent with the exception of HRZM and BPa (ranked 4th and 11th, respectively, based on T_10,_ and 7th and 9th, respectively, based on T_50_).

**TABLE 5.**
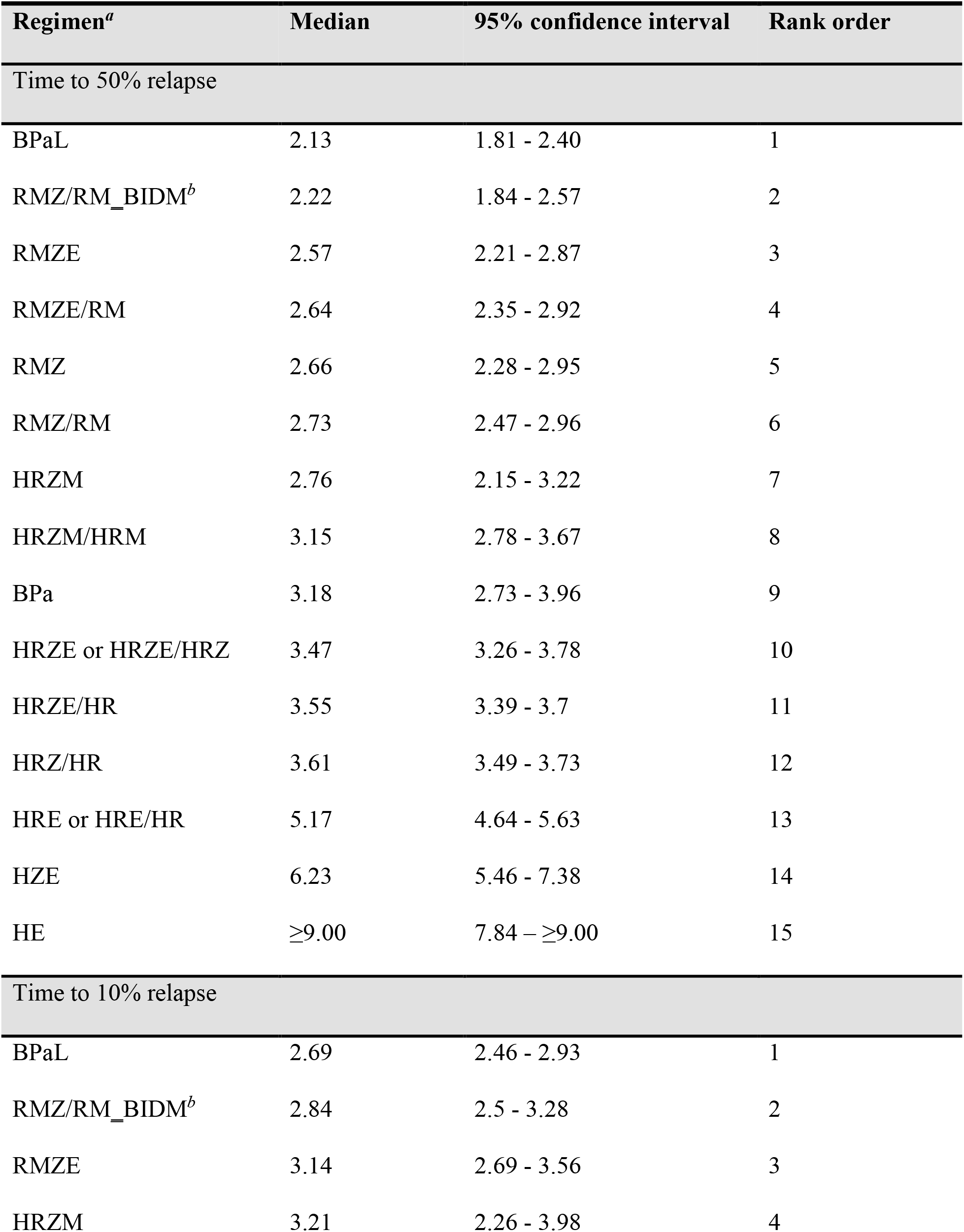
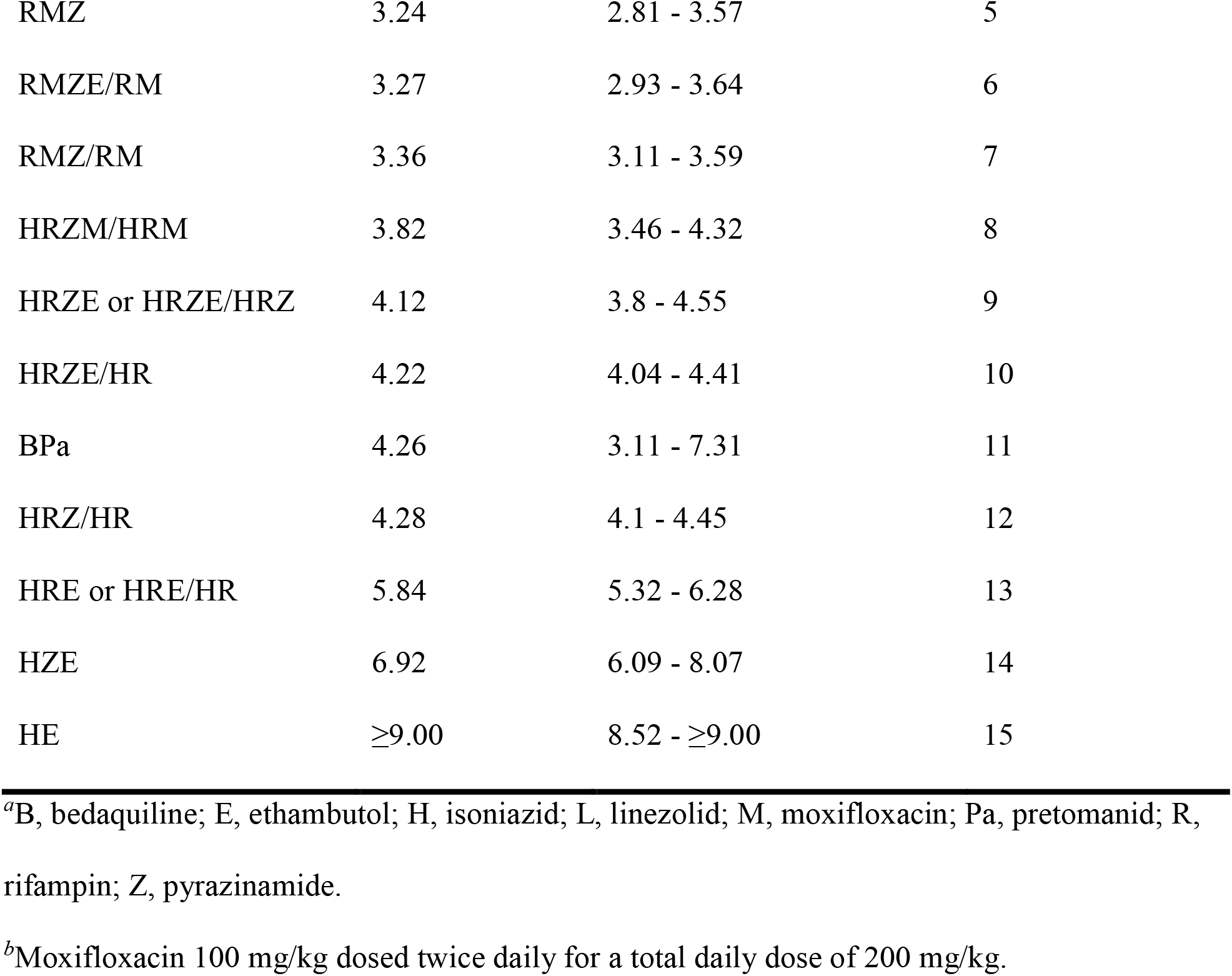
Regimen-specific T_50_ and T_10_ estimates in months and corresponding rank-order

| Regimen<sup>a</sup> | Median | 95% confidence interval | Rank order |
| --- | --- | --- | --- |
| Time to 50% relapse |  |  |  |
| BPaL | 2.13 | 1.81 - 2.40 | 1 |
| RMZ/RM_BIDM<sup>b</sup> | 2.22 | 1.84 - 2.57 | 2 |
| RMZE | 2.57 | 2.21 - 2.87 | 3 |
| RMZE/RM | 2.64 | 2.35 - 2.92 | 4 |
| RMZ | 2.66 | 2.28 - 2.95 | 5 |
| RMZ/RM | 2.73 | 2.47 - 2.96 | 6 |
| HRZM | 2.76 | 2.15 - 3.22 | 7 |
| HRZM/HRM | 3.15 | 2.78 - 3.67 | 8 |
| BPa | 3.18 | 2.73 - 3.96 | 9 |
| HRZE or HRZE/HRZ | 3.47 | 3.26 - 3.78 | 10 |
| HRZE/HR | 3.55 | 3.39 - 3.7 | 11 |
| HRZ/HR | 3.61 | 3.49 - 3.73 | 12 |
| HRE or HRE/HR | 5.17 | 4.64 - 5.63 | 13 |
| HZE | 6.23 | 5.46 - 7.38 | 14 |
| HE | ≥9.00 | 7.84 – ≥9.00 | 15 |
| Time to 10% relapse |  |  |  |
| BPaL | 2.69 | 2.46 - 2.93 | 1 |
| RMZ/RM_BIDM<sup>b</sup> | 2.84 | 2.5 - 3.28 | 2 |
| RMZE | 3.14 | 2.69 - 3.56 | 3 |
| HRZM | 3.21 | 2.26 - 3.98 | 4 |
| RMZ | 3.24 | 2.81 - 3.57 | 5 |
| RMZE/RM | 3.27 | 2.93 - 3.64 | 6 |
| RMZ/RM | 3.36 | 3.11 - 3.59 | 7 |
| HRZM/HRM | 3.82 | 3.46 - 4.32 | 8 |
| HRZE or HRZE/HRZ | 4.12 | 3.8 - 4.55 | 9 |
| HRZE/HR | 4.22 | 4.04 - 4.41 | 10 |
| BP<sub>a</sub> | 4.26 | 3.11 - 7.31 | 11 |
| HRZ/HR | 4.28 | 4.1 - 4.45 | 12 |
| HRE or HRE/HR | 5.84 | 5.32 - 6.28 | 13 |
| HZE | 6.92 | 6.09 - 8.07 | 14 |
| HE | ≥9.00 | 8.52 - ≥9.00 | 15 |

- <sup>a</sup>B, bedaquiline; E, ethambutol; H, isoniazid; L, linezolid; M, moxifloxacin; Pa, pretomanid; R, rifampin; Z, pyrazinamide. <sup>b</sup>Moxifloxacin 100 mg/kg dosed twice daily for a total daily dose of 200 mg/kg.

**FIG 3.**
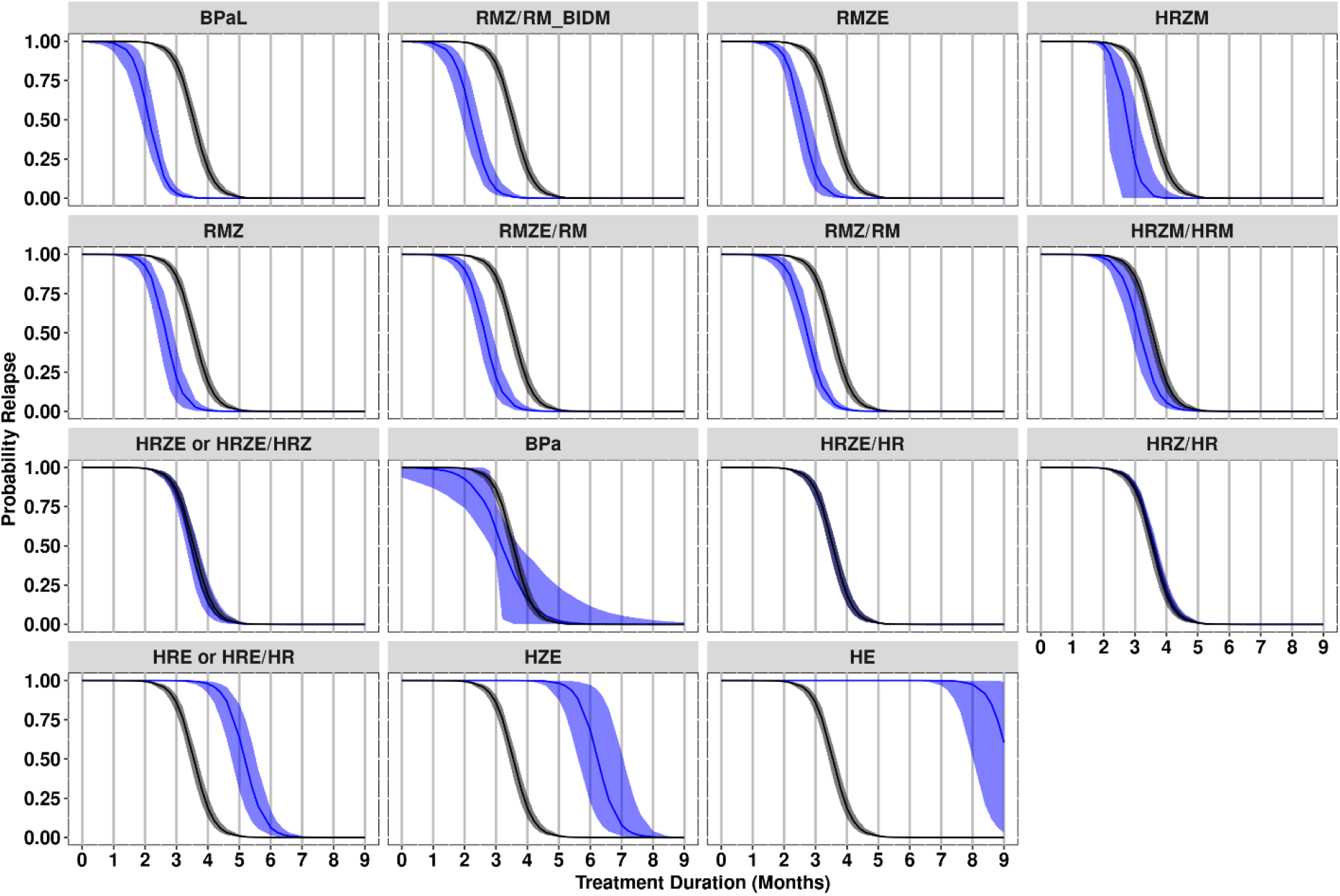
Relapse probability versus treatment duration by regimen. Blue lines and areas represent the median and 95% confidence interval, respectively, of relapse probability versus treatment duration profiles for each regimen as derived by bootstrap (N = 500 runs). Black lines and areas represent HRZE/HR as the clinical standard of care regimen. For comparative purposes, all regimens are presented as normalized to the median covariate values. “RMZ/RM_BIDM” denotes a version of the RMZ/RM regimen where moxifloxacin was administered at 100 mg/kg twice daily for a total daily dose of 200 mg/kg. E, ethambutol; H, isoniazid; M, moxifloxacin; R, rifampin; Z, pyrazinamide.

**FIG 4.**
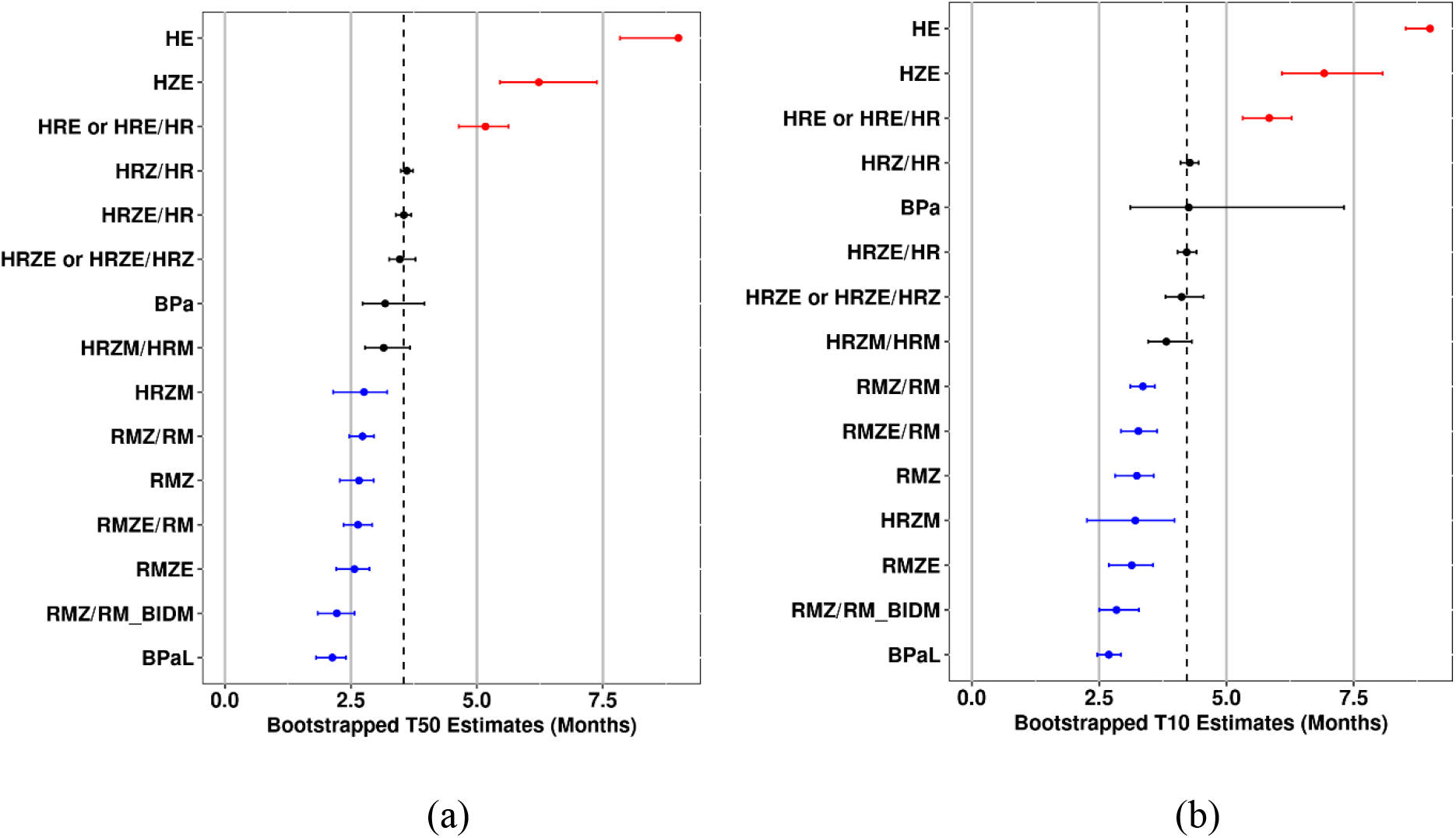
Forest plots of T_50_ and T_10_ relapse probability by regimen. Median estimates and 95% confidence intervals from bootstrapped datasets (n = 500 runs) for a) T_50_ and b) T_10_ metrics. Regimens are in descending rank orders for metrics based on median value, with red and blue coloring indicating regimens for which the confidence intervals are completely above or below the median value for HRZE/HR (included as the clinical standard of care regimen). For comparative purposes, all regimens are presented as normalized to the median covariate values. “RMZ/RM_BIDM” denotes a version of the RMZ/RM regimen where moxifloxacin was administered at 100 mg/kg twice daily for a total daily dose of 200 mg/kg. E, ethambutol; H, isoniazid; M, moxifloxacin; R, rifampin; Z, pyrazinamide.

## DISCUSSION

Historically, results from RMM studies have been reported in raw data tables with limited supporting statistical analyses limiting comparisons, interpretation, and decision-making across RMM studies and regimens. Through the application of model-based meta-analysis approaches to a large dataset of 28 studies, we have been able to improve the understanding and interpretability of regimen performance in the relapsing mouse model of TB infection. Model-based analyses have a distinct advantage over standard statistical methods used to analyze RMM studies in that the underlying mathematical model allows researchers to compare regimens not solely on the basis of *<u>proportions</u>* of mice in a treatment group relapsing at pre-specified treatment durations, but rather to obtain the relapse *<u>probability</u>* at any treatment duration (within the approximate range of durations experimentally explored). We accomplished this in the present study through a relatively simple logistic regression approach, which utilized observed binary relapse data for individual mice to estimate the INT and SLP parameters that describe the logit-linear relationship between relapse probability and treatment duration for each regimen. It is noted that this is analogous to that applied previously by Mourik, et al. (29), as well as more recently by Mudde et al. (30), which utilized a sigmoid maximum effect (E_max_) model. Although based on slightly different assumptions, both mathematical models enable the calculation of model-based parameters of interest, as well as the derivation of continuous relapse probability versus treatment duration profiles as seen in Fig. 3. Such model-based outputs are highly informative when interpreting regimen performance in RMM studies and enable quantitative comparisons to control regimens based on metrics such as the T_10_ that are analogous to clinically relevant endpoints of interest (i.e., treatment duration required to achieve an acceptably low cure rate).

Aside from the similarity in the underlying mathematical models, the present analysis extends beyond that of models based on single relapse studies (29, 30). Given the significantly larger dataset of more than 1500 mice from 28 studies, the present model-based meta-analysis methodology was able to account for inter-study covariate and random effects on the INT and SLP parameters. An advantage of this mixed-effects modeling approach was that it partitioned the observed variability in the data into inter-study variability and residual variability, which enabled quantification of inter-study standard deviations for INT and SLP as well as exploration of study-level variables as possible sources of the observed inter-study variability. The two significant covariates identified during model development as being significant contributors to inter-study variability, inoculum amount (INOC) and baseline lung bacterial burden (BASE), accounted for more than 37% and 23% of the observed inter-study variability in INT and SLP, respectively. The effect of each covariate on the overall relapse probability versus treatment duration profile is illustrated in Fig. 5 for HRZE/HR under the assumption of no interaction between covariate effects and treatment. In both cases, as the covariate value increases, the curve shifts to the right, thereby increasing the probability of relapse at a given treatment duration. This is intuitive, as it has been demonstrated that administration of a higher inoculum of *Mtb* to BALB/c mice results in a greater bacterial burden in the lung, a more severe infection, and a longer treatment duration required to prevent relapse (31). Given the correlation between inoculum size and bacterial burden at treatment start (Pearson correlation = 0.46), the joint effect estimated for HRZE/HR across the observed combinations of these variables ranges from 2.68 to 4.58 months for T_50_ and 3.41 to 5.74 months for T_10_. Hence, it is clear that the potential influence on study results is significant, and highlights the importance of accounting for these inter-study sources of variability when considering both the design (i.e., controlling the inoculum size) as well as in the analysis and interpretation of RMM study data.

**FIG 5.**
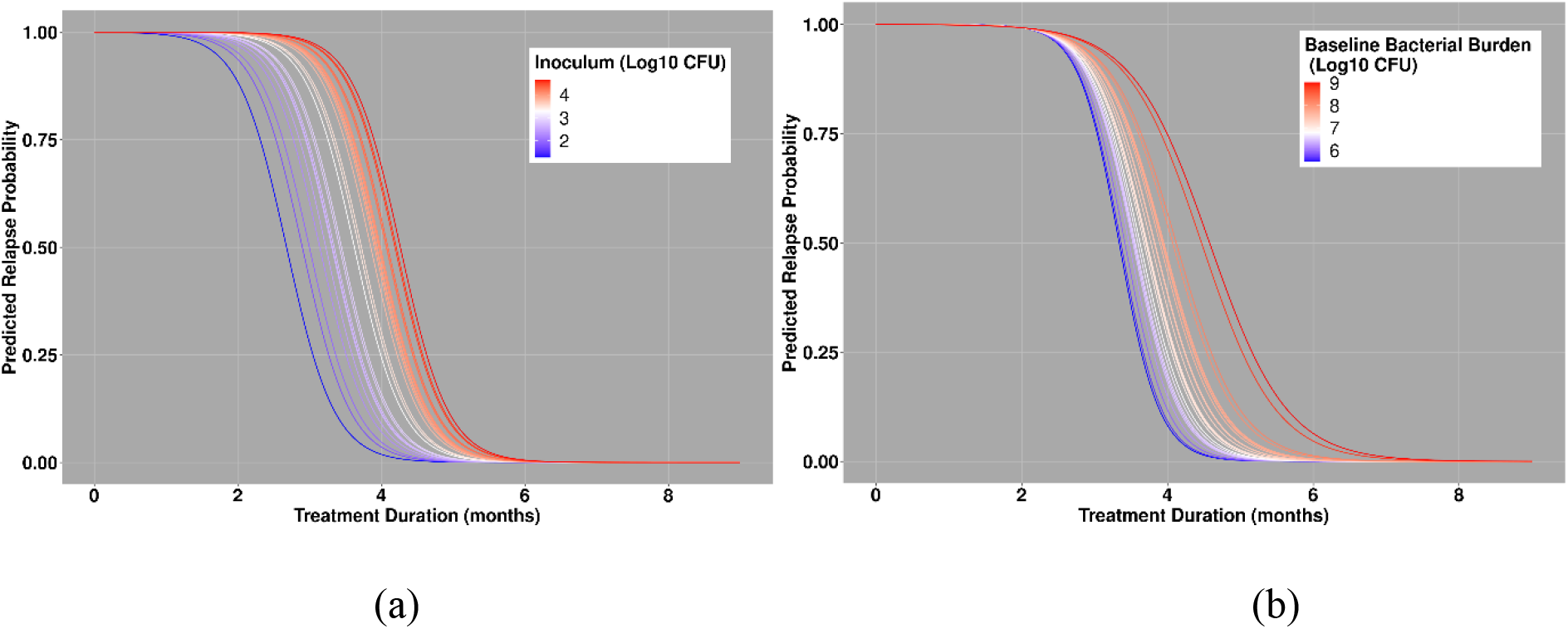
Simulations of model with range of covariate effects. Simulations of covariate effects from range of data for a) inoculum (log_10_ CFU/mL) and b) baseline bacterial burden (log_10_ CFU/mL).

The ability to account for study-level covariates and quantify inter-study variability highlights a further advantage to the model-based meta-analysis methodology applied herein, in that unbiased regimen-specific parameters were estimated simultaneously across data from all studies. Specifically, the treatment regimen fixed-effects parameters correspond to the covariate-normalized relapse probability versus treatment duration profiles after adjustment for residual inter-study variability. This is significant in that it allows for robust “apples-to-apples” comparisons of all regimens in the dataset, despite many not being evaluated together in the same experiment. That is important in light of the T_50_ estimates presented elsewhere for certain regimens (29, 30), as the impact of the significant covariates identified herein as well as other influential covariates that may not be known at this time, must be considered for cross-study comparisons. In this regard it is noted that the ability to make cross-study comparisons within this meta-analysis were bolstered by the presence of reference regimens (e.g., HRZ/HR), which helped to partition inter-study variability from treatment effects, especially for those regimens for which data were available from only a single study or where the number of treatment durations was limited. In the latter case, even with a reference regimen “anchor,” model stability often necessitated grouping similar regimens together to share an identical fixed-effect term for either INT or SLP. Groupings are indicated in Table 4, although it is noteworthy that multiple regimens did not share the same fixed effects for both parameters except for HRE and HRE/HR as well as HRZE and HRZE/HRZ. Hence, aside from these regimens which are treated as identical (which is reasonable based upon their similarity, being differentiated only on the presence and absence of ethambutol beyond two months treatment duration), the structure of the final model enabled the calculation of regimen-specific relapse probability versus treatment duration profiles.

With regard to relative regimen efficacy, the results in Fig. 3, Fig. 4, and Table 5 illustrate that, after adjusting for covariate values and inter-study variability, trends in regimen efficacy follow expected patterns. For example, the established significance of rifampin and/or pyrazinamide in shortening the treatment duration required to prevent relapse with the standard of care HRZE regimen is clearly seen by the stepwise decreases in the median T_10_ value from HE > HZE > HRE > HRZE. Similarly, the lack of differentiation between HRZ/HR and HRZE/HR supports the generally accepted premise that ethambutol adds little to the efficacy of the HRZ-based regimen in the BALB/c RMM. The treatment shortening effect of fluoroquinolone-based regimens is also clearly seen, with median T_10_ values between 0.40 months (HRZM/HRM) and 1.38 months (RMZ/RM [twice-daily 100 mg/kg moxifloxacin]) shorter than HRZE/HR. This is notable in that the treatment-shortening effect seen in initial/early mouse studies was a consideration in advancing fluoroquinolone-containing RZ-based regimens into clinical trials with a four-month treatment duration as compared to the six-month treatment duration for the standard of care HRZE/HR regimen (32, 33). However, even under the optimistic assumption of a direct 1-to-1 translation of relative regimen performance from mice to humans, the efficacy estimates obtained from our model suggest that fluoroquinolone-containing RZ-based regimens may only be capable of shortening treatment duration by approximately one month, consistent with the conclusions reached by Li, et al., as well as Wallis, et al. (15, 34). These findings are also in line with the position outlined by Lanoix, et al., that the failures seen in Phase 3 trials of four-month fluoroquinolone-containing RZ-based regimens do not reflect poor predictive performance of the RMM but rather an overly optimistic translation of RMM findings for these regimens to the clinic (35). It should be noted, however, that the treatment-shortening effect of fluoroquinolones is dependent upon the overall drug combination administered, as the combination of moxifloxacin, rifapentine, isoniazid, and pyrazinamide has recently been reported as an effective four-month regimen (36). Overall, our findings demonstrate that model-based analysis of RMM data provides results that are not only consistent with previous studies, but also builds on the understanding and interpretability of RMM studies by providing robust, quantitative, and meaningful measures of relative regimen efficacy.

Although the model developed in the current study is limited to only those regimens included in the historical CPTR dataset, the model itself is not “static” and allows for continuous updating to include data from new studies and regimens and may be applied to inform the design of future RMM studies (through *in silico* trial simulation). In the latter, further iterations of the model are being developed using emerging data on new regimens to assess the contribution of various regimen components and help select promising regimens for future study. An example of this approach is described in this report where the model was updated through incorporation of additional data for the novel BPa and BPaL regimens. The model-based estimates show that the novel two-drug BPa backbone is as efficacious in the RMM as the standard of care regimen and that the addition of linezolid as a third component provides significant improvement in regimen efficacy (Table 5), with the resulting BPaL regimen clearly being superior to HRZE/HR. Of importance, though, the model-based estimates indicate that BPaL may be no better than fluoroquinolone-based regimens at shortening overall treatment duration. Consequently, while BPaL is a highly efficacious regimen, it may not be capable of meeting the goal of treatment shortening to <two months. Prioritization of new regimens should be informed using the present model-based approach through accumulation of new RMM data on new candidate regimens pooled with the existing historic dataset, and subsequent model re-estimation. Multiple RMM studies are currently planned or ongoing that will generate additional data, with the design of several studies directly informed using the current model. Specifically, the mixed-effects logistic regression model has been applied to undertake RMM trial simulations to evaluate attributes such as number of mice per arm, number of treatment durations, inoculum size, different hypothetical treatment regimens, and inclusion of control arms. The relapse outcomes of simulated “virtual” mice are then analyzed using the same model-based approach to assess the performance of the various designs in terms of overall bias and precision on T_50_ and T_10_. This approach, which has been highly instrumental in selecting and refining the design of RMM studies to increase precision while minimizing the number of mice required for each study, will be detailed in a subsequent report.

Taken together, the model-based meta-analysis presented herein represents an improvement in the analysis, understanding, and interpretability of data from relapsing mouse model studies. By adjusting for key study-level differences and accounting for inter-study variability, this approach generates robust, quantitative, and relevant metrics of interest, such as T_50_ and T_10_, respectively, that enhance the understanding and interpretation of RMM study data and ultimately support decision making with regard to regimen selection and prioritization.

## ACKNOWLEDGMENTS

We would like to acknowledge the contributions of the Preclinical Sciences Working Group of the Critical Path to TB Drug Regimens Initiative for the critical discussions that helped to define the scope of this project, as well as for their review of the analysis plan and feedback on preliminary results. This work was supported by the Bill and Melinda Gates Foundation, which provided funding to the Critical Path to TB Drug Regimens Initiative and Cognigen Corporation.

## Supplementary Material

**TABLE S-1.**
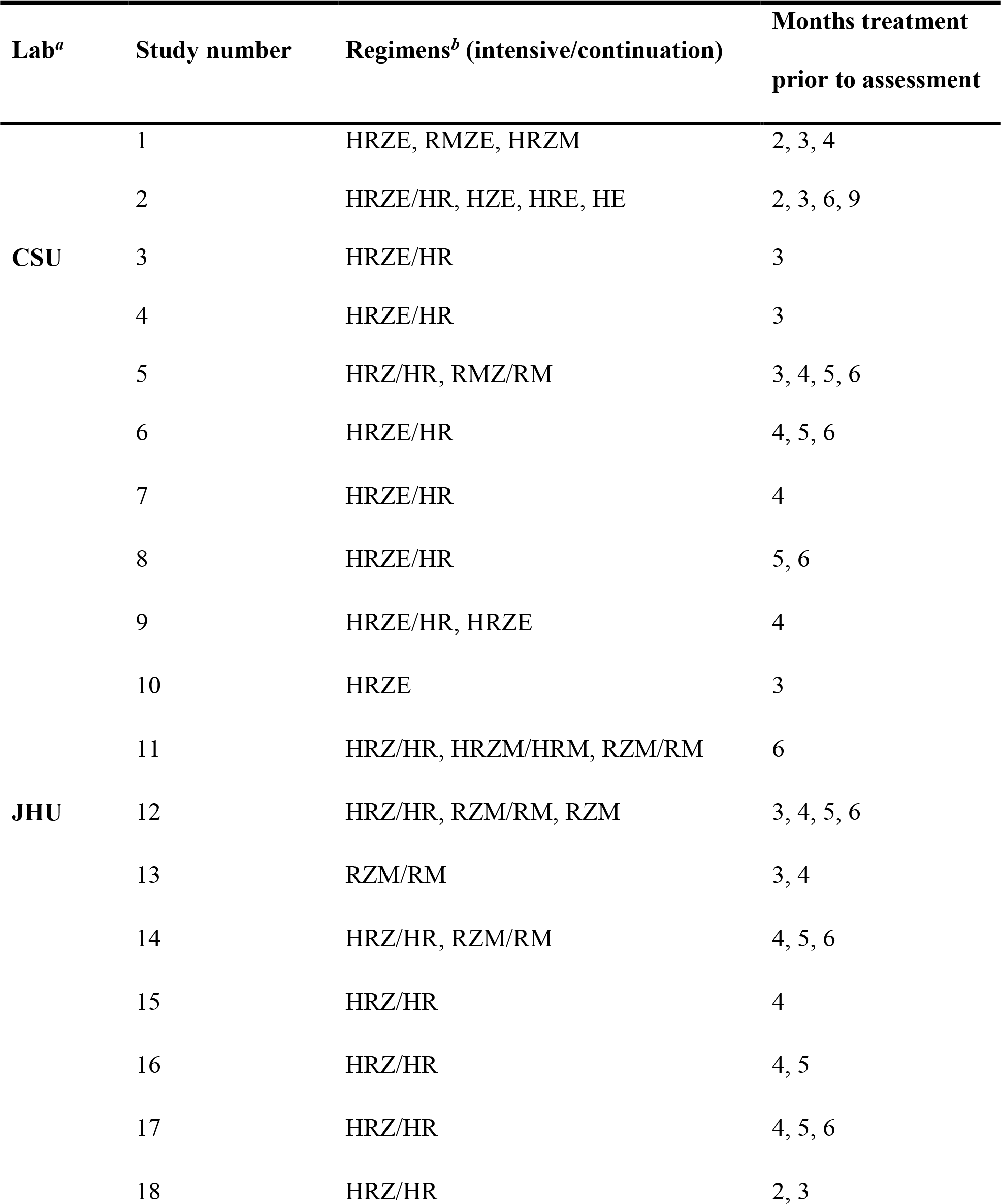
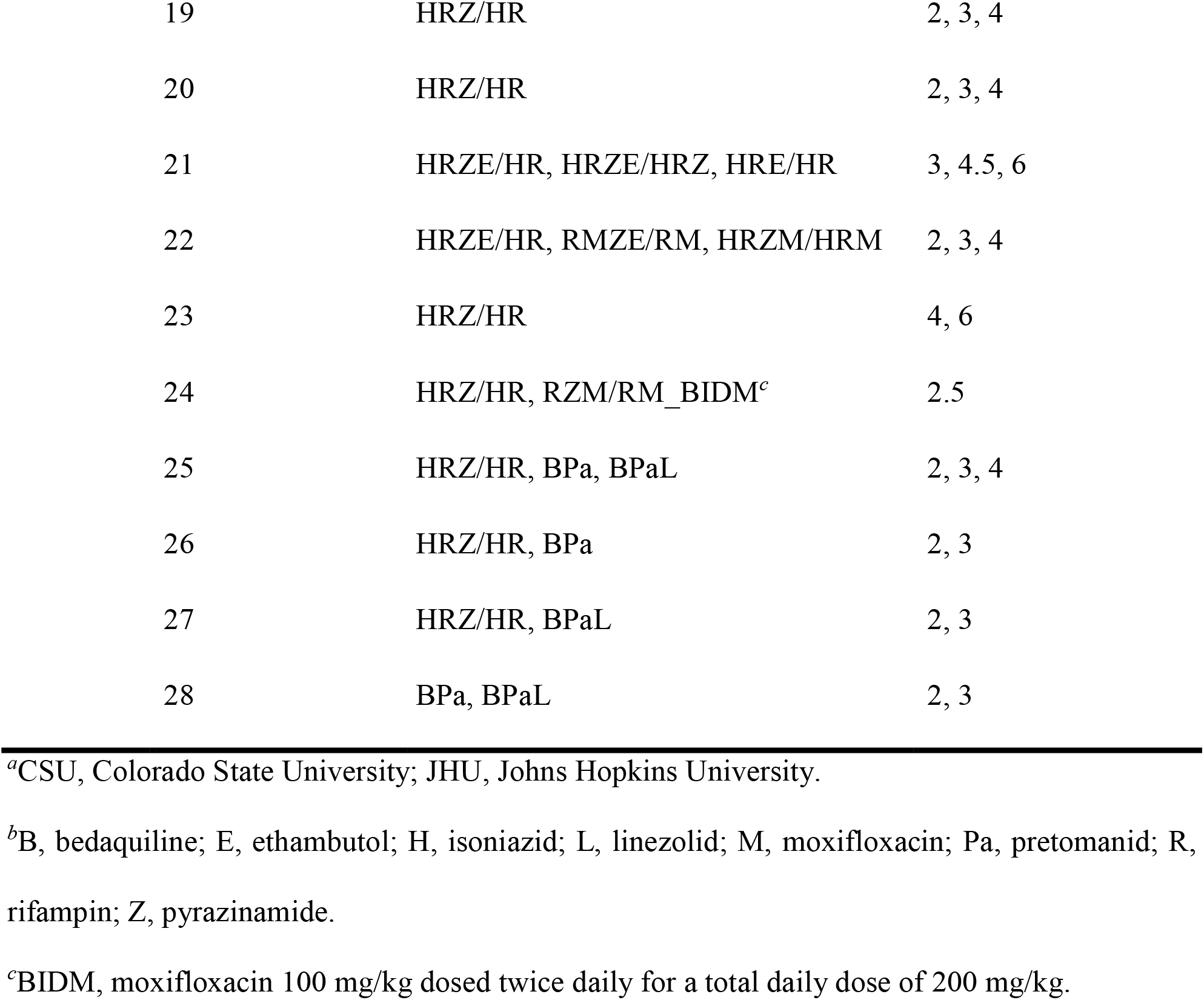
Study information by contributing lab

**FIG S-1.**
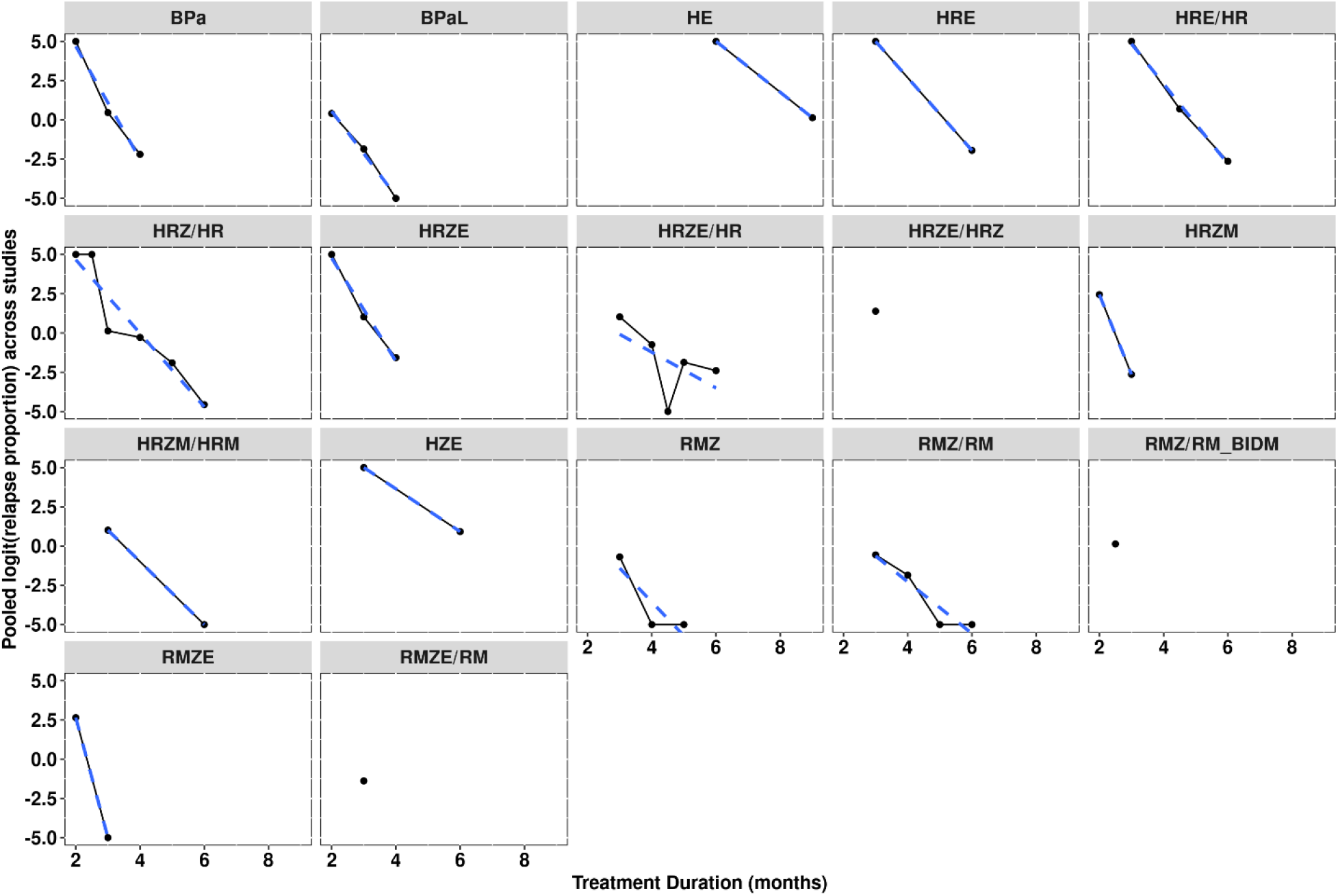
Logit-transformed relapse proportion versus treatment duration by regimen. Black dots and solid lines represent the observed relapse proportion calculated across all studies. Blue dashed lines represent a linear regression fit. “RMZ/RM_BIDM” denotes a version of the RMZ/RM regimen where moxifloxacin was administered at 100 mg/kg twice daily for a total daily dose of 200 mg/kg. M, moxifloxacin; R, rifampin; Z, pyrazinamide.

**FIG S-2.**
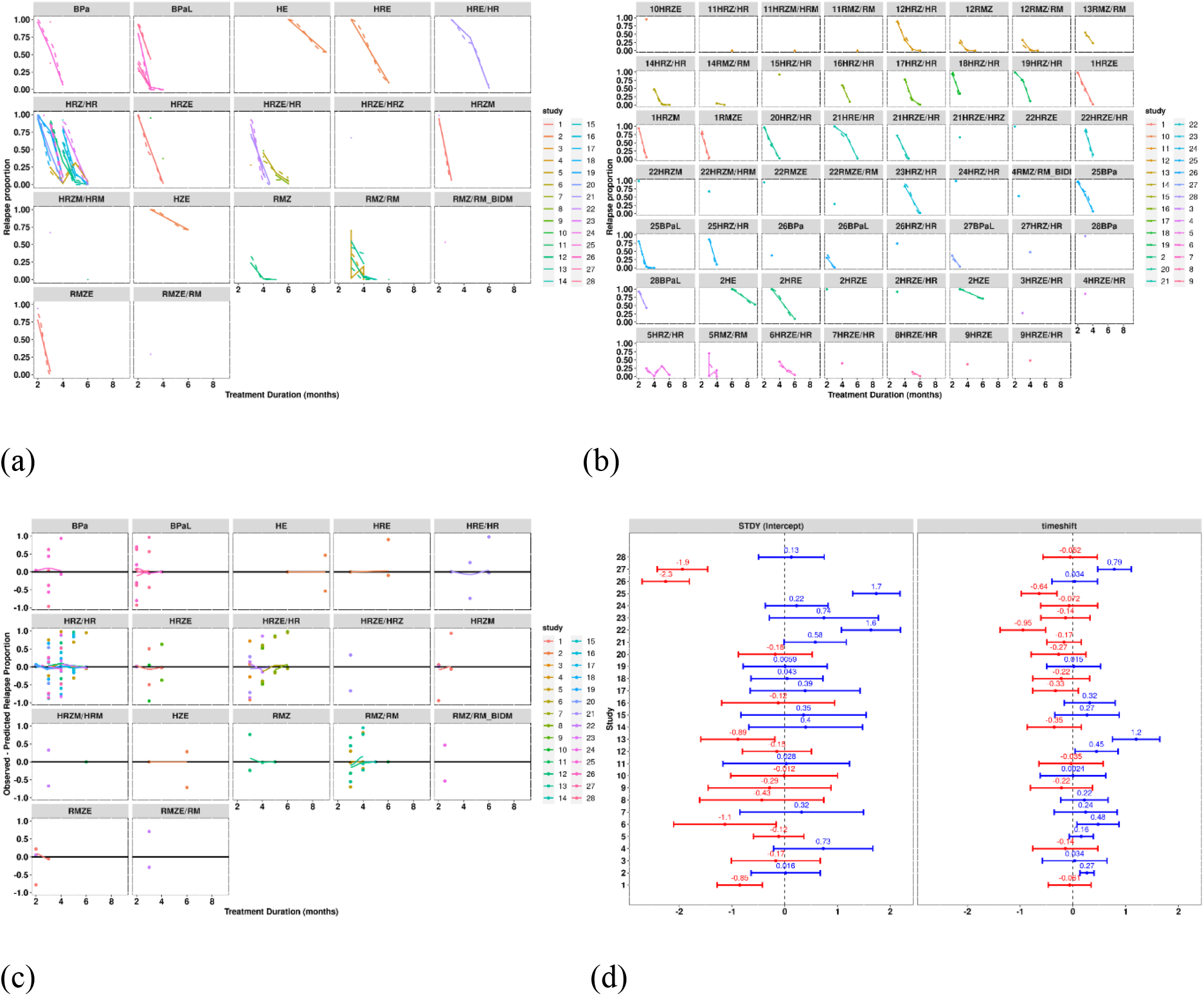
Goodness-of-fit plots for the final model. Goodness-of-fit plots for the final model: a) relapse proportion (observed [points] with overlaid predictions [solid lines]) versus time stratified by regimen, b) relapse proportion (observed [points] with overlaid predictions [solid lines]) versus time stratified by study and regimen, c) observed minus predicted relapse proportion versus time stratified by regimen and study, and d) forest plot of random effects on intercept (INT) and slope (SLP). Overall, the model is able to describe the data well. “RMZ/RM_BIDM” denotes a version of the RMZ/RM regimen where moxifloxacin was administered at 100 mg/kg twice daily for a total daily dose of 200 mg/kg. M, moxifloxacin; R, rifampin; Z, pyrazinamide.

**FIG S-3.**
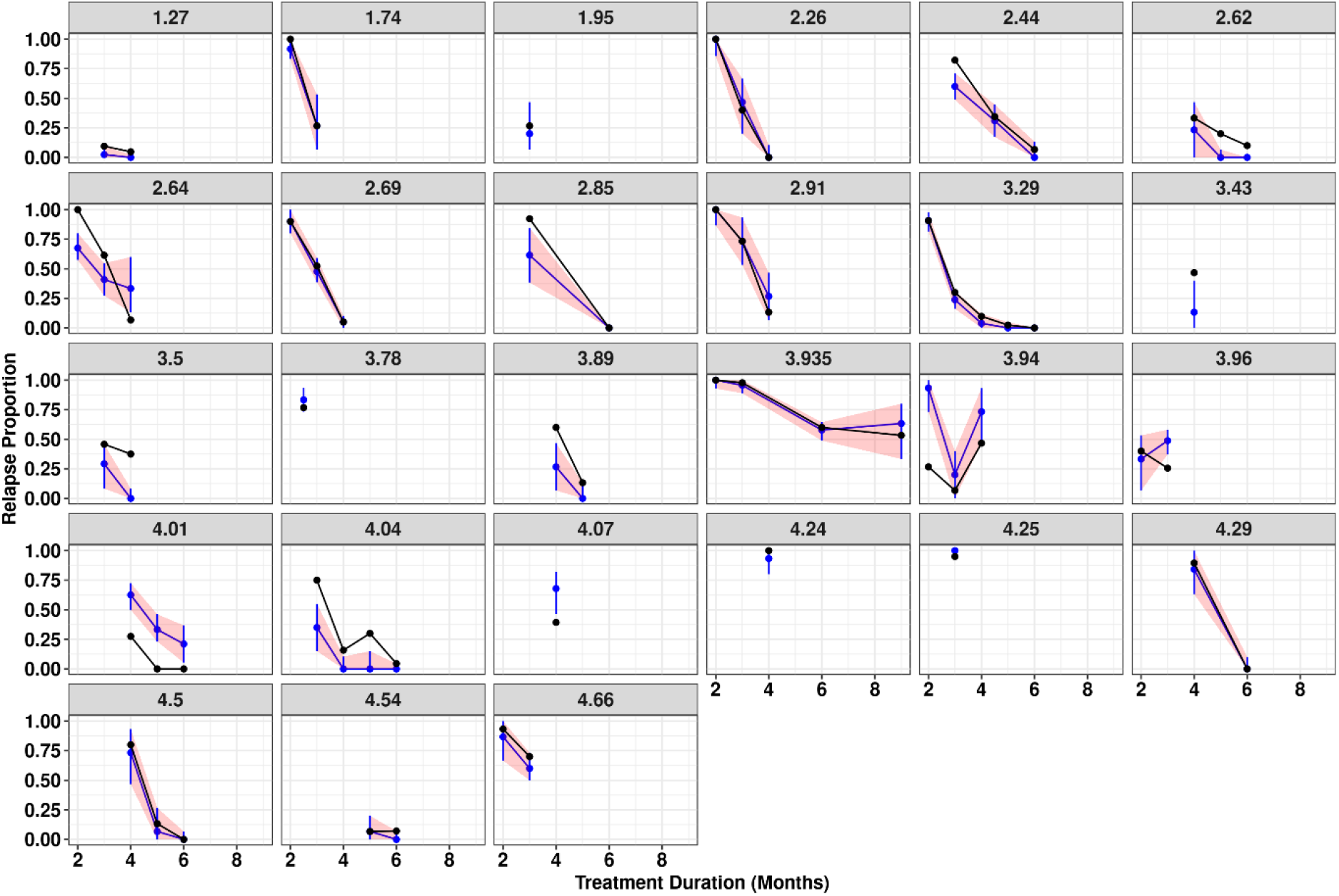
Visual predictive check for the final model – stratified by inoculum (Log_10_ CFU). Black dots and solid lines represent the observed relapse proportion. Blue dots and solid lines represent the median prediction from the final model. The red shaded area represents the 90% prediction interval.

**FIG S-4.**
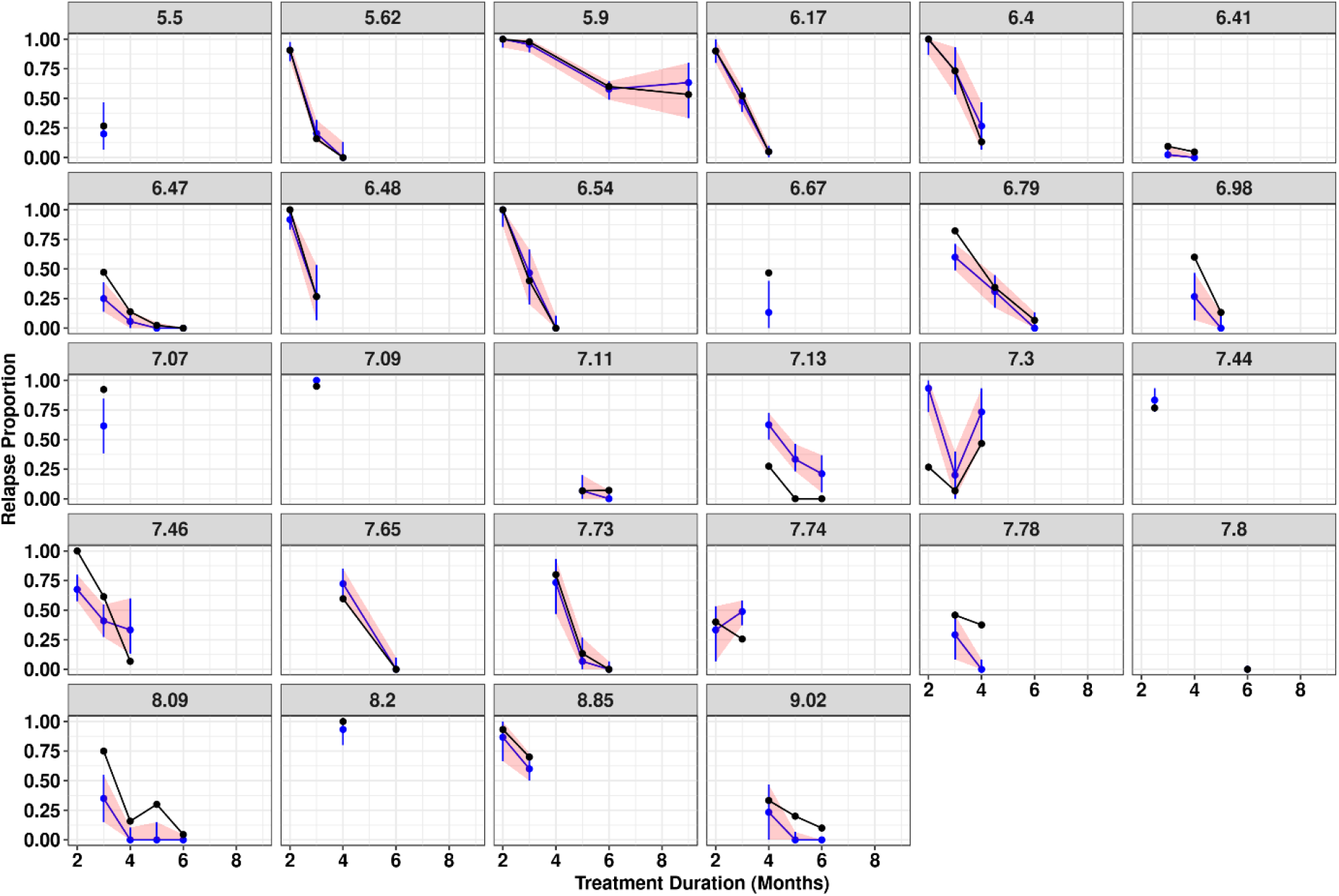
Visual predictive check for final model – stratified by baseline bacterial burden (Log_10_ CFU). Black dots and solid lines represent the observed relapse proportion. Blue dots and solid lines represent the median prediction from the final model. The red shaded area represents the 90% prediction interval.

**FIG S-5.**
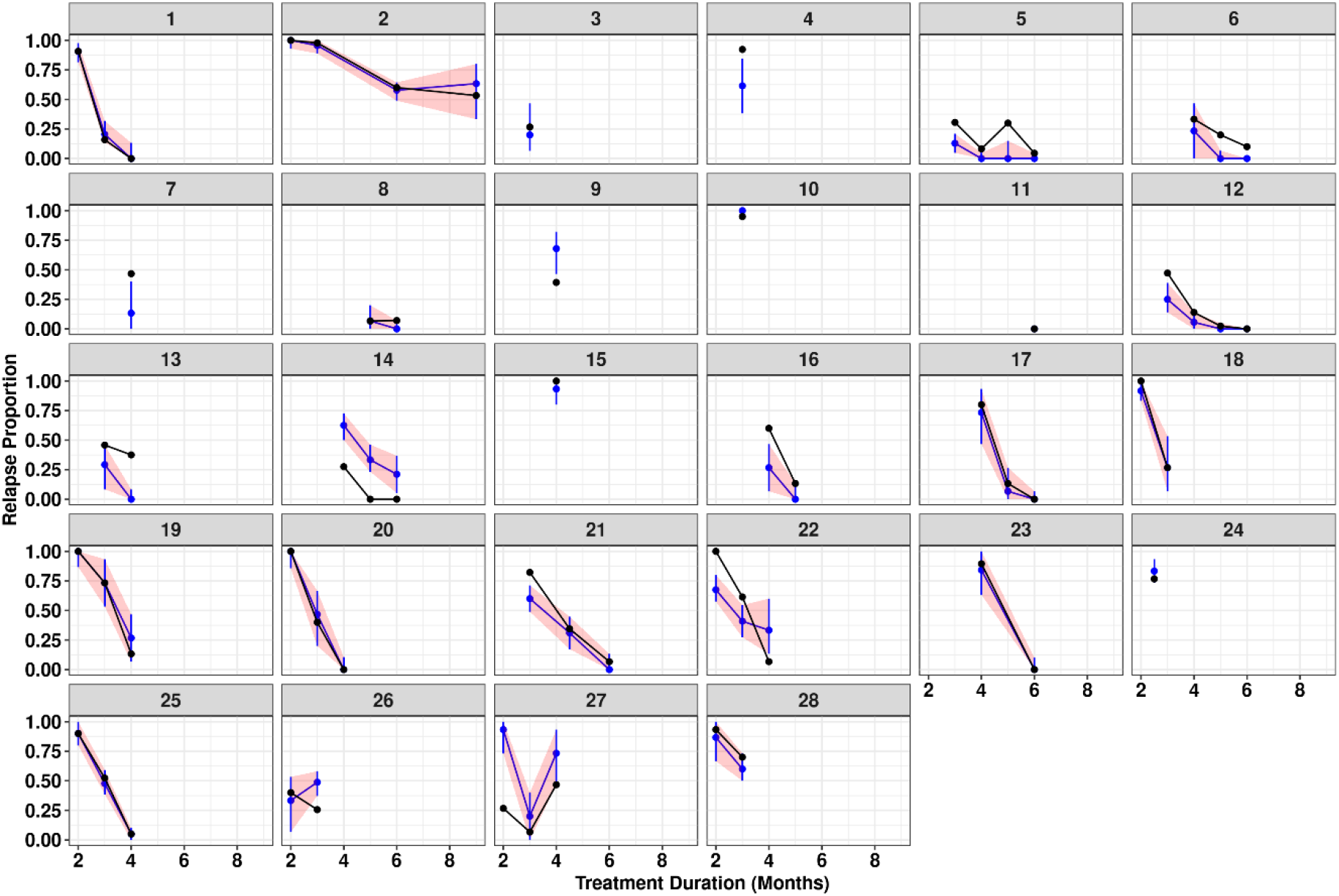
Visual predictive check for final model – stratified by study. Black dots and solid lines represent the observed relapse proportion. Blue dots and solid lines represent the median prediction from the final model. The red shaded area represents the 90% prediction interval.

## References

1. Harding E. 2020. WHO global progress report on tuberculosis elimination. Lancet Respir Med 8(1):19. doi: 10.1016/S2213-2600(19)30418-7. Epub 2019 Nov 6. Erratum in: Lancet Respir Med. 15 Nov 2019; PMID: 31706931.

2. World Health Organization. 2018. Latent TB Infection: Updated and consolidated guidelines for programmatic management. http://www.who.int/tb/publications/2018/latent-tuberculosis-infection/en.

3. World Health Organization, Global Tuberculosis Programme. 2020. Global tuberculosis report. https://www.who.int/publications/i/item/9789240013131.

4. European Medicines Agency. 21 Nov 2013. Committee for Medicinal Products for Human Use. Summary of Opinion: Deltyba. https://www.ema.europa.eu/en/documents/smop-initial/chmp-summary-positive-opinion-deltyba_en.pdf.

5. Food and Drug Administration. 2019. Pretomanid product insert. Silver Spring (MD). https://www.accessdata.fda.gov/drugsatfda_docs/label/2019/212862s000lbl.pdf.

6. Food and Drug Administration. 2012. Sirturo (bedaquiline) product insert. Silver Spring (MD). http://www.accessdata.fda.gov/drugsatfda_docs/label/2012/204384s000lbl.pdf.

7. Combs DL, O’Brien RJ, Geiter LJ. 1990. USPHS Tuberculosis Short-Course Chemotherapy Trial 21: effectiveness, toxicity, and acceptability. The report of final results. Ann Intern Med 112(6):397–406.

8. Critical Path Institute. Critical path to TB drug regimens. Available from: https://c-path.org/programs/cptr/.

9. Imperial MZ, Nahid P, Phillips PPJ, Davies GR, Fielding K, Hanna D, Hermann D, Wallis RS, Johnson JL, Lienhardt C, Savic RM. 2018. A patient-level pooled analysis of treatment-shortening regimens for drug-susceptible pulmonary tuberculosis. Nat Med 2018;24(11):1708–1715. doi: 10.1038/s41591-018-0224-2. Epub 2018 Nov 5. Erratum in: Nat Med. 2019 Jan;25(1):190. PMID: 30397355; PMCID: PMC6685538.

10. Gumbo T, Lenaerts AJ, Hanna D, Romero K, Nuermberger E. 2015. Nonclinical models for antituberculosis drug development: a landscape analysis. J Infect Dis 211 Suppl 3:S83–95.

11. Franzblau SG, DeGroote MA, Cho SH, Andries K, Nuermberger E, Orme IM, Mdluli K, Angulo-Barturen I, Dick T, Dartois V, Lenaerts AJ. 2012. Comprehensive analysis of methods used for the evaluation of compounds against Mycobacterium tuberculosis. Tuberculosis (Edinb) 92(6):453–88. doi: 10.1016/j.tube.2012.07.003. Epub 2012 Aug 30. PMID: 22940006.

12. De Groote MA, Gruppo V, Woolhiser LK, Orme IM, Gilliland JC, Lenaerts AJ. 2012. Importance of confirming in vivo efficacy data of novel antibacterial drug regimens against various strains of Mycobacterium tuberculosis. Antimicrob Agents Chemother 56(2):731–738.

13. De Groote MA, Gilliland JC, Wells CL, Brooks EJ, Woolhiser LK, Gruppo V, Peloquin CA, Orme IM, Lenaerts AJ. 2011. Comparative studies evaluating mouse models used for efficacy testing of experimental drugs against M. tuberculosis. Antimicrob Agents Chemother 55(3):1237–1247.

14. Lenaerts AJ, Chapman PL, Orme IM. 2004. Statistical limitations to the Cornell model of latent tuberculosis infection for the study of relapse rates. Tuberculosis (Edinb) 84(6):361–364.

15. Li SY, Irwin SM, Converse PJ, Mdluli KE, Lenaerts AJ, Nuermberger EL. 2015. Evaluation of moxifloxacin-containing regimens in pathologically distinct murine tuberculosis models. Antimicrob Agents Chemother 59(7):4026–4030.

16. Saini V, Ammerman NC, Chang YS, Tasneen R, Chaisson RE, Jain S, Nuermberger E, Grosset JH. 24 May 2019. Treatment-shortening effect of a novel regimen combining clofazimine and high-dose rifapentine in pathologically distinct mouse models of tuberculosis. Antimicrob Agents Chemother 63(6):e00388–19. https://doi:10.1128/AAC.00388-19.

17. Ammerman NC, Swanson RV, Bautista EM, Almeida DV, Saini V, Omansen TF, Guo H, Chang YS, Li SY, Tapley A, Tasneen R, Tyagi S, Betoudji F, Moodley C, Ngcobo B, Pillay L, Bester LA, Singh SD, Chaisson RE, Nuermberger E, Grosset JH. 26 Jun 2018. Impact of clofazimine dosing on treatment shortening of the first-line regimen in a mouse model of tuberculosis. Antimicrob Agents Chemother 62(7):e00636–18. https://doi:10.1128/AAC.00636-18.

18. Tyagi S, Ammerman NC, Li S-Y, Adamson J, Converse PJ, Swanson RV, Almeida DV, Grosset JH. 20 Jan 2015. Clofazimine shortens the duration of the first-line treatment regimen for experimental chemotherapy of tuberculosis. Proc Natl Acad Sci U S A 112(3):869-74. https://doi:10.1073/pnas.1416951112.

19. Rosenthal IM, Zhang M, Williams KN, Peloquin CA, Tyagi S, Vernon AA, Bishai WR, Chaisson RE, Grosset JH, Nuermberger EL. Dec 2007. Daily dosing of rifapentine cures tuberculosis in three months or less in the murine model. PLoS Med 4(12):e344. https://doi:10.1371/journal.pmed.0040344.

20. Lanoix J-P, Betoudji F, Nuermberger E. 7 Dec 2015. Sterilizing activity of pyrazinamide in combination with first-line drugs in a C3HeB/FeJ mouse model of tuberculosis. Antimicrob Agents Chemother 60(2):1091–1096. https://doi:10.1128/AAC.02637-15.

21. Tasneen R, Li S-Y, Peloquin CA, Taylor D, Williams KN, Andries K, Mdluli KE, Nuermberger EL. 2011. Sterilizing activity of novel TMC207- and PA-824-containing regimens in a murine model of tuberculosis. Antimicrob Agents Chemother 55(12):5485–5492. https://doi:10.1128/AAC.05293-11.

22. Nuermberger E, Tyagi S, Tasneen R, Williams KN, Almeida D, Rosenthal I, Grosset JH. 2008. Powerful bactericidal and sterilizing activity of a regimen containing PA-824, moxifloxacin, and pyrazinamide in a murine model of tuberculosis. Antimicrob Agents Chemother 52(4):1522–1524. https://doi:10.1128/AAC.00074-08.

23. Tasneen R, Betoudji F, Tyagi S, Li S-Y, Williams K, Converse PJ, Dartois V, Yang T, Mendel CM, Mdluli KE, Nuermberger EL. 26 Oct 2015. Contribution of oxazolidinones to the efficacy of novel regimens containing bedaquiline and pretomanid in a mouse model of tuberculosis. Antimicrob Agents Chemother 60(1):270–277. doi: 10.1128/AAC.01691-15. PMID: 26503656; PMCID: PMC4704221.

24. Akaike H. 1973. Information theory and an extension of the maximum likelihood principle, p 267–281. In Petrov BN, Csáki F (ed), 2nd International Symposium on Information Theory, Tsahkadsor, Armenia, USSR, September 2-8, 1971, Budapest: Akadémiai Kiadó. Republished in Kotz S, Johnson NL, eds. (1992), Breakthroughs in Statistics, I, Springer-Verlag, pp. 610-24.

25. R Core Team. 2017. R: a language and environment for statistical computing. R Vienna, Austria: Foundation for Statistical Computing. https://www.R-project.org/.

26. RStudio Team. 2020. RStudio: Integrated Development for R. RStudio, PBC, Boston, MA. http://www.rstudio.com/.

27. Dardis C. 2015. LogisticDx: Diagnostic tests for models with a binomial response. R package Version 0.2. https://CRAN.R-project.org/package=LogisticDx.

28. Wickham H. 2016. ggplot2: Elegant Graphics for Data Analysis. 2nd ed. New York (NY): Springer. 276 p.

29. Mourik BC, Svensson RJ, de Knegt GJ, Bax HI, Verbon A, Simonsson USH, de Steenwinkel JEM. 9 Apr 2018. Improving treatment outcome assessment in a mouse tuberculosis model. Sci Rep 8(1):5714. doi: 10.1038/s41598-018-24067-x. PMID: 29632372; PMCID: PMC5890284.

30. Mudde SE, Alsoud RA, van der Meijden A, Upton AM, Lotlikar MU, Simonsson USH, Bax HI, de Steenwinkel JEM. 19 Feb 2021. Predictive modeling to study the treatment-shortening potential of novel tuberculosis drug regimens, toward bundling of preclinical data. J Infect Dis jiab101. https://doi:10.1093/infdis/jiab101.

31. Nuermberger EL. 2017. Preclinical efficacy testing of new drug candidates. Microbiol Spectr 5(3) doi: 10.1128/microbiolspec. TBTB2-0034-2017. PMID: 28643624.

32. Nuermberger EL, Yoshimatsu T, Tyagi S, Williams K, Rosenthal I, O’Brien RJ, Vernon AA, Chaisson RE, Bishai WR, Grosset JH. 2004. Moxifloxacin-containing regimens of reduced duration produce a stable cure in murine tuberculosis. Am J Respir Crit Care Med 170(10):1131–1134.

33. Nuermberger EL, Yoshimatsu T, Tyagi S, O’Brien RJ, Vernon AN, Chaisson RE, Bishai WR, Grosset JH. 2004. Moxifloxacin-containing regimen greatly reduces time to culture conversion in murine tuberculosis. Am J Respir Crit Care Med 169(3):421–426.

34. Wallis RS, Wang C, Meyer D, Thomas N. 5 Aug 2013. Month 2 culture status and treatment duration as predictors of tuberculosis relapse risk in a meta-regression model. PLoS One. https://doi.org/10.1371/journal.pone.0071116.

35. Lanoix JP, Chaisson RE, Nuermberger EL. 2016. Shortening tuberculosis treatment with fluoroquinolones: lost in translation? Clin Infect Dis 62(4):484–490.

36. Dorman SE, Nahid P, Kurbatova EV, Phillips PPJ, Bryant K, Dooley KE. 6 May 2021. Four-month rifapentine regimens with or without moxifloxacin for tuberculosis. N Engl J Med 384(18):1705–1718. https://doi:10.1056/NEJMoa2033400.

